# Evolution of HIV virulence in response to widespread scale up of antiretroviral therapy: a modeling study

**DOI:** 10.1101/039560

**Authors:** Joshua Herbeck, John E. Mittler, Geoffrey S. Gottlieb, Steven Goodreau, James T. Murphy, Anne Cori, Michael Pickles, Christophe Fraser

**Affiliations:** Department of Global Health, University of Washington, Seattle, WA 98104; Department of Microbiology, University of Washington, Seattle, WA 98195; Departments of Medicine and Global Health, University of Washington, Seattle, WA 98195; Department of Anthropology, University of Washington, Seattle, WA 98195; Department of Infectious Disease Epidemiology, Imperial College London, London W2 1PG, UK

## Abstract

There are global increases in the use of HIV antiretroviral therapy (ART), guided by clinical benefits of early ART initiation, and the efficacy of treatment as prevention of transmission. Separately, it has been shown theoretically and empirically that HIV virulence can evolve over time. Observed virulence levels may reflect an adaptive balance between infected lifespan and per-contact transmission rate. Critically, the potential effects of widespread ART on HIV virulence are unknown. To predict these effects, we used an agent-based stochastic model to simulate changes in HIV virulence, using set point viral load as a virulence proxy. We calibrated our epidemic model to the prevalence and incidence trends of South Africa. We repeated our analysis using a separate calibration, as a sensitivity analysis, and found that predicted impact of ART on virulence was relatively insensitive to calibration. We explored two ART scenarios, at increasing coverage : ART initiation based on patients reaching a CD4 cell count threshold, or based on time elapsed since infection (a scenario that mimics the “universal testing and treatment” aspirations). We found that HIV virulence is generally unchanged in scenarios of CD4-based initiation. However, with ART initiation based on time since infection, virulence can increase moderately within several years of ART rollout, under high coverage and early treatment initiation, albeit within the context of rapidly shrinking epidemics. Our modeling study suggests that increasing virulence driven by universal test and treat is likely not a major public health concern, but could be monitored in sentinel surveillance, in a manner similar to transmitted drug resistance.

## INTRODUCTION

Worldwide, 15 million HIV-infected individuals are receiving ART (1). To take advantage of the benefits of earlier treatment initiation for both individual endpoints (better disease prognosis (2)) and population endpoints (decreased rates of onward transmission (3)), UNAIDS has set ambitious targets of 90-90-90 by 2020. (90% of all people living with HIV will know their HIV status, 90% of all people with diagnosed HIV infection will receive sustained antiretroviral therapy, and 90% of all people receiving antiretroviral therapy will have viral suppression) (http://www.unaids.org/sites/default/files/media_asset/90-90-90_en_0.pdf).

However, it is unknown how widespread scale up of ART may affect HIV virulence. HIV virulence, defined here as the rate of disease progression in untreated infections, is commonly estimated via the proxies of set point viral load (SPVL; the viral load after acute infection but prior to AIDS), baseline CD4+ T-cell count, rate of CD4+ T-cell decline, immune activation, or viral replicative capacity.

Modeling studies of HIV virulence have suggested that, in epidemics where ART use is not widespread, HIV may adaptively evolve toward an intermediate level of virulence, to balance the per-contact transmission rate with the infected lifespan (4-6). These findings are consistent with the “trade-off” theory of virulence evolution, where an optimal virulence level should exist that maximizes the (pathogen’s) lifetime transmission success (7, 8). For example, low HIV virulence (low SPVL) will result in decreased per-contact infectivity but more lifetime transmissions in untreated infections (due to longer infected lifespans); high virulence (high SPVL) will result in higher per-contact infectivity but fewer total transmissions (due to shorter infected lifespans) (4). These HIV virulence models have focused on SPVL, as SPVL is prognostic for the rate of disease progression (9-12). Importantly, SPVL is a phenotype with the necessary requirements for adaptive evolution: variation in the host population (13, 14); correlation with the per-contact transmission rate (viral fitness) (15-19); and at least partially determined by genotype (*i.e.* heritable across transmission pairs) (20-24).

Empirical studies of HIV virulence have revealed unexplained variation in population-level estimates of virulence and trends in virulence among local HIV epidemics (local as defined by cohort, population, or country) (14). Hypothesized explanations for this variation include: founding viral lineages of local epidemics had different virulence levels (6); different epidemic ages have allowed for varying amounts of HIV adaptation to the host population (6, 25); virulence estimates have been affected by sampling biases related to improvements in HIV screening and diagnosis (6) or the time between last negative and first positive HIV antibody tests (which can affect the precision of SPVL measurements) (14); and, related to the current study, differences in the history and extent of antiretroviral therapy (ART) in each local epidemic (25).

Large-scale “treatment as prevention” or “universal testing and treatment” programs will likely shift the distribution of HIV transmissions by individual stage of infection (26), potentially modifying the balance between per-contact transmission rate and the length of the infected lifespan. It is possible that biomedical interventions that extend the lifespans of HIV-infected individuals will shorten the (effective) viral lifespans and result in increased HIV virulence: less virulent viruses will no longer reap the benefits of longer infected lifespans. Alternatively, it is possible that individuals infected with the most virulent viruses will initiate treatment the earliest, thus providing an added evolutionary advantage to less virulent viruses. In short, viruses that transmit the most before treatment initiation may have a selective advantage. However, because of the range of possible interactions and counter-acting selection pressures, these evolutionary scenarios are hard to intuit, and so need to be studied within the context of a mathematical model. It is important to identify specific ART-related scenarios in which HIV virulence could evolve, and how rapidly virulence could change.

Our goal was to predict, using a stochastic, agent-based modeling approach, the effect of ART scale-up on HIV virulence. We used SPVL as a virulence proxy to investigate whether ART can apply evolutionary pressure on the balance between HIV per-contact infectivity and the infected lifespan. Additionally, we examined whether different scale-up scenarios (*e.g.*, based on observed clinical practice or national treatment guidelines) for ART eligibility and initiation mediate this pressure. Our model allowed for fine-scale variation among viral lineages in virulence (individual SPVL), and allowed for population-based levels of virulence (mean SPVL) to change over time. We used our model to predict changes in HIV incidence and prevalence that result from ART intervention programs (which are comparable to earlier modeling studies), and concurrently to predict the changes in virulence in these same epidemic scenarios.

## RESULTS

### Calibration

Our primary results focus on the calibration of a stochastic, agent-based model (6) to the HIV epidemic of South Africa (Table 1). We chose to calibrate the model in this way first, because the epidemic in South Africa is one of the largest, globally, and secondly, to make our study explicitly and readily comparable to the 12 models of ART and HIV incidence that were documented and compared by Eaton *et al.* (27). These 12 models, and comparisons among them represent the fullest attempt, to date, at quantifying the potential impact of ART on HIV incidence in a high-incidence generalized epidemic. These 12 models included simulations starting in 1990 with ART rollout starting in 2012. The models were compared based on epidemiological endpoints measured at 8 and 38 years after ART rollout (2020 and 2050, respectively); we followed this same structure. Our simulations were initiated with 2% HIV prevalence and initial mean SPVL of 3.5, 4.5 or 5.5 log_10_ copies/mL. Incidence (per 100 person years) rose quickly to ∼3%, followed by a decline to ∼1.75% and stabilization around year 2010 (Figure 1). Prevalence rose to ∼12%, followed by a decline to ∼8%. 20% of transmissions occured in the from individuals in their first year of infection; the majority of transmissions occured in chronic infection. These epidemic outputs, in the absence of ART, were similar to outputs from the 12 models compared by Eaton *et al.* (27).

**Figure 1.**
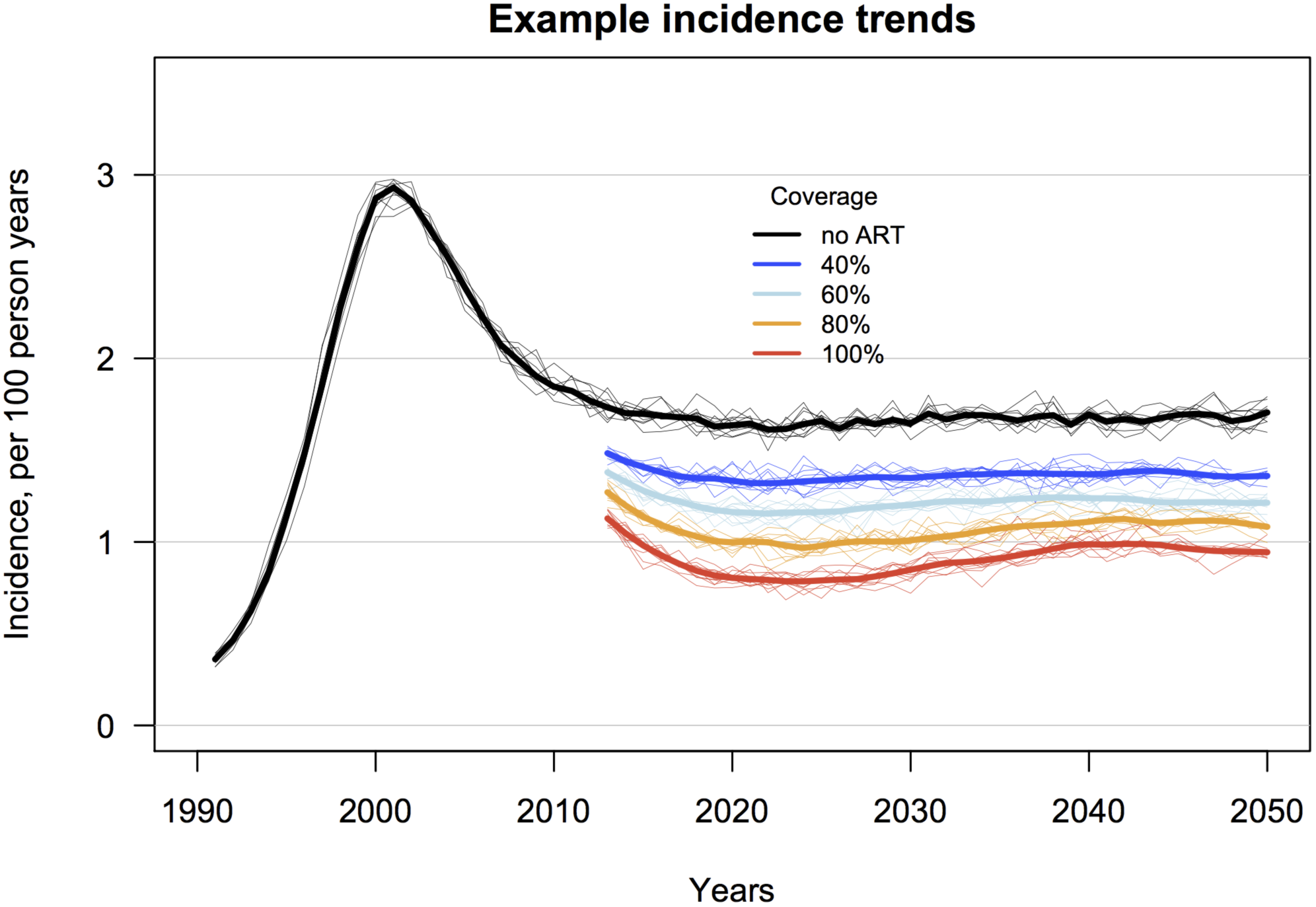
Simulated trends in HIV incidence for ART scenarios of 40, 60, 80 and 100% coverage (individual probability of receiving ART with complete adherence) and CD4 count threshold for treatment initiation <350 cells/mL, versus the counterfactual epidemic simulation with no ART. Shown are LOESS regression lines for ten random replicates for each ART coverage scenario (thin lines), and the mean of these replicates (thick lines). Initial mean SPVL was 4.5 log_10_ copies/mL.

**Table 1.** Parameters of the model and initial values. Parameters with an asterisk had different values between the **original** and **alternate** model calibrations.

| Parameter | Value |
| --- | --- |
| Demographic and behavioral |  |
| Initial overall population size | $1 \times 10^5$ individuals |
| Initial number of infected | $2 \times 10^3$ individuals |
| Minimum relationship duration | 0.1 years |
| Maximum relationship duration | 5.0 years |
| Subgroups defined by relationship duration* | original: <0.5 yrs, 0.5-2.5 yrs, >2.5 yrs<br>alternate: 1 group (no subgroups) |
| Probability of sexual contact, in each group* | original: 1.0, 0.05, 0.03 per day<br>alternate: 1.0 |
| Mean degree* | original: 0.9<br>alternate: 0.7 |
| Virologic |  |
| Viral load at time zero | 10 copies/mL |
| Viral load at peak viremia | $1.0 \times 10^7$ copies/mL (15, 35) |
| Time to peak viremia | 21 days (15, 35) |
| Total time of acute infection | 91 days (15) |
| Viral load progression rate, natural log | 0.05 per year (37) |
| Viral load at AIDS (CD4<200) | $5.0 \times 10^6$ copies/mL (37) |
| Set point viral load |  |
| Variance of $\log_{10}$ SPVL | 0.7 (13) |
| Heritability of SPVL across transmissions ($h^2$) | 0.36 (24) |
| Mutational variance | 0.20 |
| Transmission |  |
| Maximum transmission rate | 0.005 per day (4, 5) |
| Viral load at 0.5 max transmission rate | 13,938 copies/mL (4, 5) |
| Hill coefficient, transmission function | 1.02 (4, 5) |
| Shape parameter, transmission function | 3.46 (4, 5) |
| Disease progression |  |
| See Cori *et al.* for CD4 wait time matrix stratified by SPVL (38) |  |
| Antiretroviral therapy |  |
| Time to initiation after becoming eligible by CD4 | 1 year (27) |
| Viral load after ART initiation | 50 copies/mL |

### Virulence evolution in the absence of ART

Starting with initial SPVL distributions that reflected low, intermediate, and high virulence (mean SPVLs of 3.5, 4.5, and 5.5 log_10_ copies/mL, respectively), HIV evolved toward an intermediate level of virulence (Figure S1). Additionally, as predicted under stabilizing selection, the population variance of SPVL decreased over time in all scenarios. Under the current model assumptions, the evolutionary optimal mean SPVL, in the absence of ART, is predicted to be ∼4.70 log_10_ RNA copies/mL. We have previously performed sensitivity analyses for the effects of viral and behavioral parameters on SPVL levels and SPVL evolution in our model (6); we summarize these here. The inferred optimal virulence (mean SPVL) was insensitive to variation in the following viral parameters: a) rate of viral load increase in chronic infection (*s* from Equation 1); b) maximum transmission rate (*B*_max_ in Equation 2); c) maximum time to AIDS; and d) peak viremia in acute infection. The rate at which mean SPVL evolved to an inferred population-level optimum (as seen in Figure S1) depended principally on *B*_max_; mean SPVL increased more rapidly in the early years of epidemics with higher *B*_max_ values, before arriving at similar mean SPVL levels (6). The inferred optimal virulence (mean SPVL) was sensitive to the mean degree of the sexual network, defined as the average number of sexual partners that each person has at any given time. Higher mean degrees were associated with higher mean SPVL. Because we were interested in comparing mean SPVL between simulations with and without ART, this sensitivity did not affect our main findings. However, our alternate model calibration, performed as a sensitivity analysis and described below, includes a lower mean degree and thus lower mean SPVL.

### Virulence evolution in the presence of ART

#### Time since infection thresholds for ART initiation, representing Universal Test and Treat

With ART initiation thresholds based on time elapsed since infection (without consideration of CD4 count or stage of infection), virulence was slightly increased by 2020 (8 years after ART rollout), except in scenarios with 100% coverage, in which mean SPVL was increased by 0.2 log_10_ with initiation at three years after infection (Figure 2). By 2050, however, increased virulence was seen at all coverage levels, at all initiation times. The largest increases were seen with ART initiation at two, three and four years after infection and ART coverage at and above 60%. In these scenarios the maximum increases in mean SPVL were ∼0.4 log_10_ copies/mL by 2050 (*e.g.* from ∼4.7 to ∼5.1 log_10_ copies/mL) (Figure 2 and Figure S2). These increases in virulence only occurred in the context of very large reductions in incidence (Figure S3). Figure 3 shows the specific example of increasing mean SPVL for ART initiation at three years after infection, for ART coverage ranging from 40% to 100%. Similar plots for SPVL trends under each of the evaluated time thresholds, at increasing coverage levels, are shown in the Supporting Information (Figure S4).

**Figure 2.**
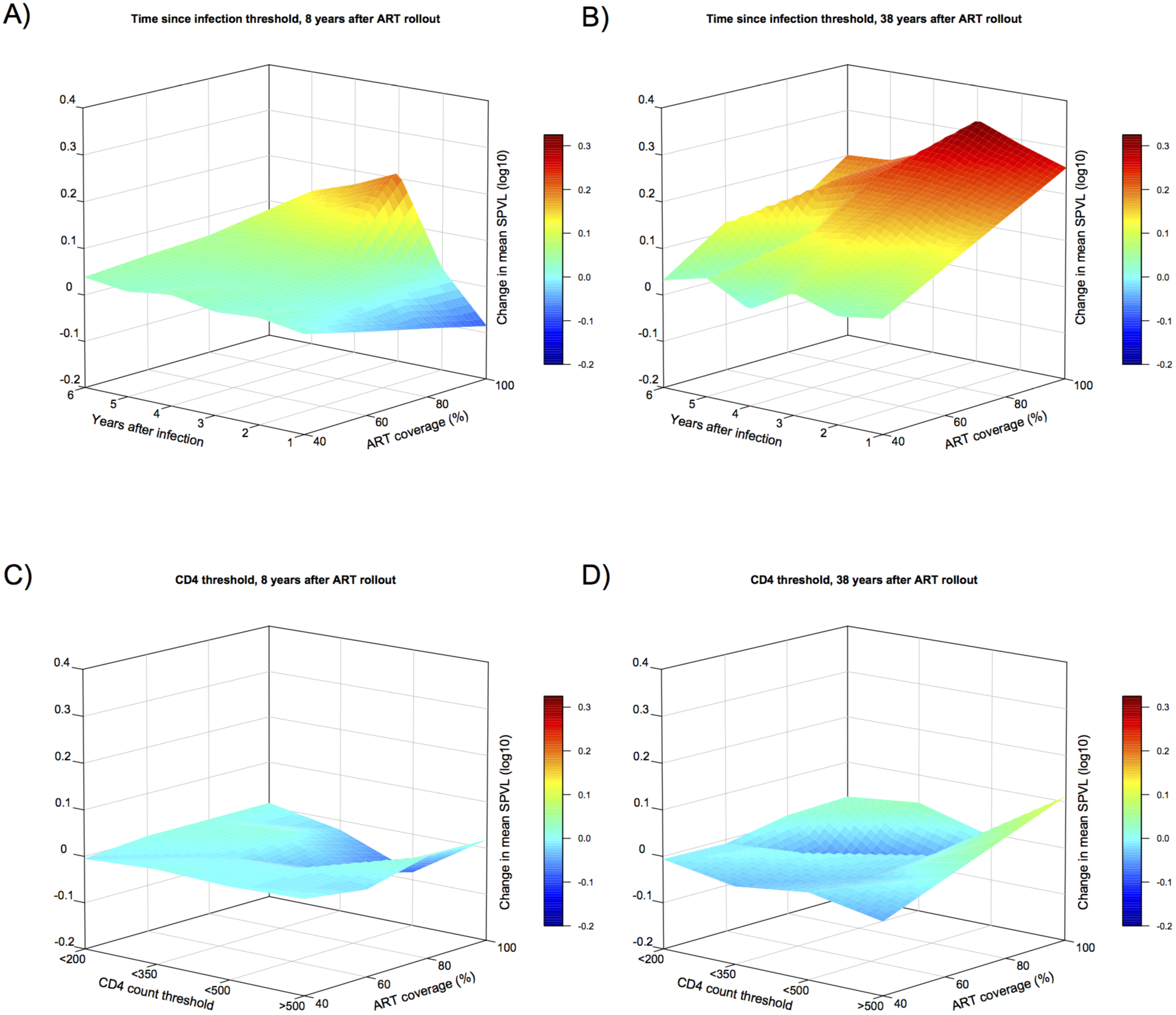
Surface plots showing change in mean SPVL between epidemic simulations with and without ART, for scenarios of increasing ART coverage (individual treatment probability) and ART initiation based either on time since infection or CD4 count threshold. For epidemic scenarios with ART initiation based on time since infection, **A)** shows mean SPVL change 8 years after ART rollout (from year 2012 to year 2020), and **B)** shows mean SPVL change 38 years after rollout (from year 2012 to year 2050). For epidemic scenarios with ART initiation based on CD4 count, **C)** and **D)** show mean SPVL at 8 and 38 years after rollout, respectively.

#### CD4 count thresholds for ART initiation

With CD4 count thresholds for ART initiation, mean SPVLs at eight and 38 years after ART rollout (2020 and 2050) were generally unchanged between ART and counterfactual (no ART) simulations (Figure 2 and Figure S5). Small decreases in virulence (decreases in mean SPVL up to ∼0.10 log_10_ copies/mL) were seen by 2020 for the CD4<500 ART initiation threshold at higher coverage levels (80% and 100%), with the magnitude of change greater with increasing coverage. These decreases in virulence were not maintained by 2050. Slight increases in virulence were seen by 2050 in the CD4>500 scenario (all individuals are eligible for ART) with high coverage; incidence was near zero in these scenarios (Figure S6). Additional plots for SPVL trends under the examined range of CD4 thresholds, at increasing coverage levels, are shown in the Supporting Information (Figure S7).

### Incidence trends in the presence of ART

Our model produced estimates of person-years of ART needed per infection averted, and reductions in incidence, that were similar to estimates described in Eaton *et al.* (27) (Table 2). In ART simulations with 80% coverage and an ART initiation threshold of CD4<350, we observed a mean percent reduction in incidence of 40.03% (s.d. 2.31) by 2020 (eight years after ART rollout) and of 36.82% (s.d. 2.92) by 2050. In these simulations, the mean person-years of ART needed per infection averted was 7.42 (s.d. 0.22) and 8.60 (s.d. 2.47) in 2020 and 2050, respectively. As expected, we observed greater declines in incidence as ART coverage or CD4 count eligibility thresholds increased (Figure S6). Reassuringly, incidence declined even in the ART scenarios that led to increasing HIV virulence.

**Table 2.** Simulated impacts of ART, comparing models with evolving HIV virulence (a distribution of individual SPVL values and allowing for evolutionary change) and static SPVL (all individuals have SPVL of 4.5 log_10_ copies/mL and individual SPVLs are identical across transmissions). Two ART scenarios are shown for evolving and static SPVL models: ART initiation and eligibility at CD4<350 cells/ul and 80% coverage; and ART initiation and eligibility at four years elapsed after infection and 80% coverage. Values are means and standard deviations for ten replicate model runs.

|  | Time since infection threshold |  | CD4 count threshold |  |
| --- | --- | --- | --- | --- |
|  | Static SPVL | Evolving SPVL | Static SPVL | Evolving SPVL |
| Percent reduction in incidence |  |  |  |  |
| year 2020 | 53.99 ±2.51 | 54.27 ±2.92 | 32.42 ±2.83 | 40.03 ±2.31 |
| year 2050 | 43.61 ±4.03 | 38.72 ±2.23 | 31.42 ±2.45 | 36.82 ±3.46 |
| Person-years of ART per infection averted |  |  |  |  |
| year 2020 | 7.52 ±0.24 | 6.08 ±0.22 | 9.98 ±0.19 | 7.42 ±0.22 |
| year 2050 | 9.39 ±1.67 | 7.59 ±0.46 | 12.80 ±1.75 | 8.60 ±2.47 |

### Incidence trends for evolving versus static SPVL

Our HIV epidemic model includes a distribution of virulence levels among individuals and allows virulence (mean SPVL) to evolve. To assess the potential effects of these model characteristics on standard epidemic output, we repeated our simulations with a single, time invariant SPVL (4.5 log_10_ copies/mL) and 100% heritability (*i.e*., all individuals have SPVL=4.5 over the course of a simulation), and compared the output of this static SPVL model to the output of the evolving SPVL model.

With a variable and evolving SPVL, the predicted benefits of ART are greater than predicted by a model with static SPVL (in an epidemic scenario of 80% ART coverage and an ART initiation threshold of CD4<350) (Table 2). With evolving SPVL, incidence reductions are greater by years 2020 and 2050 (*e.g.* ∼40% vs ∼32% reduction by year 2020), and fewer person-years of ART are required per infection averted (*e.g.* 7.42 person-years vs. 9.98 person-years at year 2020; 8.60 vs 12.80 person-years by year 2050).

### Results from the alternate model calibration

We repeated our simulation experiments with alternate parameterization of the epidemic model, as a test of the sensitivity of our primary results (above) to the overall model calibration. These alternate epidemic simulations were initiated with 2% HIV prevalence and initial mean SPVL of 4.5 log_10_ copies/mL; incidence (per 100 person years) rises at a slower rate than in our primary model, but continues rising to ∼5% incidence by the end of the epidemic simulation (Figure S8). This alternate model produced results, with respect to HIV virulence evolution, equivalent to those obtained from the primary model calibration: ART initiation based on CD4 count results in generally unchanged HIV virulence (mean SPVL relative to the no ART counterfactual), while ART initiation based on time since infections results in moderate increases in mean SPVL, in scenarios of early initiation and high coverage (Figure S9, Figure S10 (Time trends), and Figure S11 (CD4 trends)).

## DISCUSSION

HIV virulence evolution has attracted a considerable amount of recent interest (4, 14). Separately, the effects of ART on HIV transmission, incidence, and prevalence are issues of critical public health importance (3, 27). In our analysis we have used a mathematical model to jointly examine these inter-related aspects of HIV evolution and epidemiology.

### ART scenarios and virulence evolution

Based on our model, we predict that widespread ART use will, overall, have minimal effects on HIV virulence. However, specific scenarios may yield clinically significant increases in SPVL: when treatment initiation is based solely on the time elapsed since infection (rather than a CD4 count threshold), when treatment initiation occurs relatively early after infection, and when treatment population coverage is relatively high (at or above 60%). The maximum predicted increases in mean SPVL were ∼0.4 log_10_ copies/mL by 2050, 38 years after ART rollout (*e.g.* from ∼4.7 to ∼5.1 log_10_ copies/mL). In these scenarios, the beneficial impact of ART programs could be partially mitigated by the emergence of highly virulent viruses; nonetheless, the model predicts substantial and durable reductions in incidence for these scenarios.

Table 3 compares clinically relevant results of distinct ART modeling scenarios, including 80% coverage and initiation at CD4<350 threshold (used as the ART scenario for the primary model comparisons described in Eaton *et al.* (27)), and the time since infection scenario with 80% coverage and initiation threshold of three years since infection. In the time since infection scenario, which represent universal test and treat, our model predicts increases in mean SPVL of 0.27 log_10_ copies/mL by 2050 (4.72 to 4.97) (Figure 3). Following the transmission and disease progression functions of our model (see Materials and Methods and Supporting Information), an increase of this magnitude results in a ∼8% increase in annual transmission rate (infectivity) and a ∼13% decrease in time to CD4<350. This change in infectivity hinges on the shape of our viral load and transmission function (Equation 2), which assumes that infectivity plateaus at high viral loads (4). If, rather, infectivity does not plateau but rather continues to increase with higher viral loads, increases in transmission rate due to virulence evolution will be greater (28, 29); previous studies (using different underlying functions) have predicted that SPVL increases of 0.50 log_10_ will decrease the median time to AIDS by 3 years (30), and will increase the annual transmission rate by up to 37% (4, 17, 18, 28).

**Figure 3.**
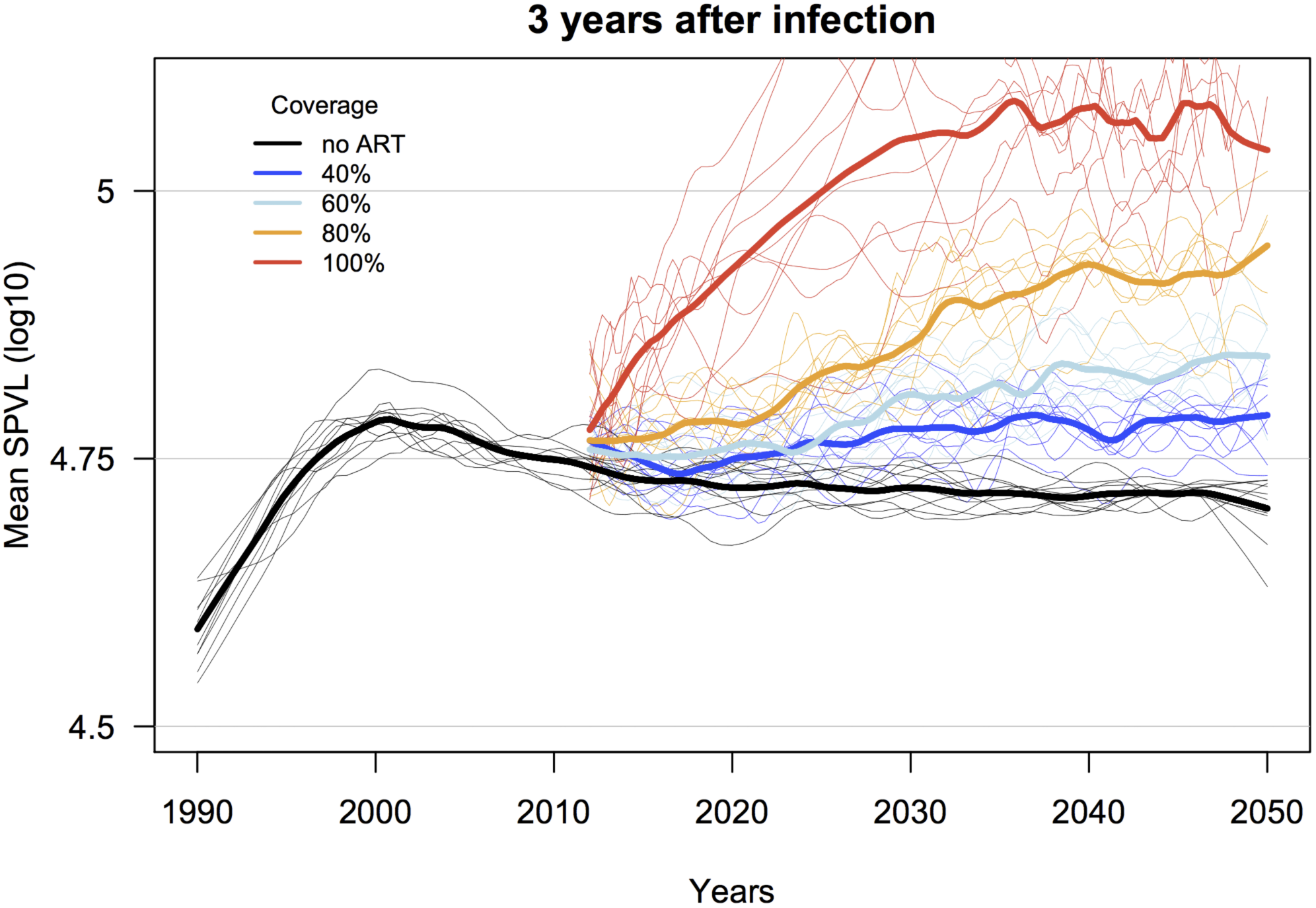
Example simulated trends in HIV virulence (mean SPVL) for scenarios of 40, 60, 80 and 100% coverage (individual probability of treatment) and **ART initiation at 3 years elapsed after infection**, versus the counterfactual simulation with no ART. Shown are LOESS regression lines for ten random replicates for each ART coverage scenario (thin lines), and the mean of these replicates (thick lines). Initial mean SPVL was 4.5 log_10_ copies/mL. ART coverage lines start at year 22, corresponding to a simulation starting at year 1990 with ART rollout at year 2012.{Bonhoeffer, 2003 #25}

**Table 3.** Impact of HIV virulence evolution given in terms of: **A)** mean SPVL; **B)** infectiousness (mean annual transmission rate); and **C)** years until specific CD4+ T-cell counts. The baseline comparison for simulated epidemics without ART is shown. ART scenarios with **80% coverage** are shown for two eligibility types: **1)** ART eligibility at CD4<350 cells/ul; and **2)** ART eligibility at four years elapsed after infection. Values are means for all new infections between 2045 and 2050 (33 to 38 years after ART, the last 5 years of 60 year epidemic runs), for 10 replicates each, for the SPVL (viral load at the end of primary infection).

|  | mean SPVL | mean annual transmission rate | mean years to CD4<500 | mean years to CD4<350 | mean years to CD4<200 |
| --- | --- | --- | --- | --- | --- |
| **No ART** | 4.72 | 0.76 | 2.40 | 4.47 | 7.03 |
| **A.** With ART; CD4<350 threshold | 4.70 | 0.75 | 2.30 | 4.27 | 6.86 |
| **B.** With ART; 3 years after infection threshold | 4.97 | 0.82 | 2.05 | 3.91 | 6.25 |
| **B.** With ART; 4 years after infection threshold | 4.90 | 0.81 | 2.18 | 4.14 | 6.73 |

The predicted dynamics of virulence evolution (Figure 2) suggest that the selective advantage of high virulence is greatest with early ART initiation times after infection (<4 years after infection). *i.e.* There is a length of infection prior to ART initiation in which the majority of high virulence viruses will have already killed their host, and thus the evolutionary cost of ART is only born by low virulence viruses. As the time threshold for ART initiation increases (up to 6 years and beyond), the observed change in mean SPVL (HIV virulence) decreases—because these scenarios are approximating the natural history of HIV without ART, and the evolutionary balance between high virulence (short lifespans with high infectivity) and low virulence (long lifespans with low infectivity) is unaffected by ART.

The generally stable virulence level predicted by the ART scenarios with CD4 count thresholds is perhaps surprising, given that the mean times to these CD4 thresholds are similar to the time thresholds for initiation that produced increased virulence (*i.e.* three to five years after infection). This can be explained by the selective advantage of higher virulence viruses being mitigated by the selective earlier removal of these same high virulence lineages (by ART), relative to lower virulence lineages. In effect, ART initiation based on CD4 count appears to balance the selective processes for and against more virulent viruses, while ART initiation based on time elapsed since infection provides a selective advantage *only* for more virulent viruses. This can be seen in the relative proportions of transmissions that occur after ART rollout, for high (defined here as >4.70 log_10_ copies/mL, the observed population average, 4.7) and low (<4.70 log_10_ copies/mL) virulence viruses (Figure S12).

### Can ART explain empirical (observed) virulence trends?

The fastest mean linear increase in SPVL (estimated after ART rollout, from 2012 to 2050) was 0.008 log_10_ copies/mL/year, for 100% coverage and ART initiation at 3 years after infection (Figure 4); 50% slower than the SPVL increase observed in the largest cohort study completed to date (0.016 log_10_ copies/mL/year; 95% CI 0.013–0.019) (31), and 40% slower than the summary trend from a meta-analysis of 8 published SPVL trends (0.013 log_10_ copies/mL/year; 95% CI -0.001–0.03) (14). Given that our maximum rate was produced in a scenario with 100% coverage, we infer that the empirical SPVL trends are likely not due to ART rollout alone— although we can not absolutely rule out that differences in the primary transmission routes of our modeled epidemic (heterosexual sex) and the empirical estimates (mostly men-who-have-sex-with-men epidemics in North America and Europe) would result in different effects of ART on virulence.

It has been suggested that decreases in HIV virulence (measured by the proxies of viral replicative capacity and viral load) observed in ART-naïve antenatal cohorts in Gaborone, Botswana (compared to Durban, South Africa) may be due partially to historical increases in ART coverage (and more substantially to the extent of viral adaptation to the host immune response) (25). Our model does not simulate the evolution of viral replicative capacity *per se*, but we can assess whether the observed difference in median viral loads (4.19 and 4.47 log_10_ copies/mL, respectively, in Gaborone and Durban) can be explained by ART. A decline of this magnitude (∼0.3 log_10_ copies/mL; *ceteris paribus*) in median VL is not seen in our model when comparing ART and counterfactual simulations; the largest decreases due to ART (median decrease of ∼0.1 log_10_ copies/mL) occurred either: A) 38 years after ART rollout, or B) at high CD4 threshold (<500) and 100% coverage within 8 years of rollout—two scenarios not consistent with the history of ART in Botswana. A recent meta-analysis estimated the mean CD4 count at ART initiation in southern Africa to be 152 cells/uL (32). Thus, the majority of HIV infected individuals are likely starting ART late in infection, regardless of a given CD4-based guideline. Our model predicts that this will result in moderate increases in HIV virulence, not decreased virulence. Qualitatively, then, we infer that ART has not contributed to the postulated decreased HIV virulence in Botswana.

## Conclusions

Our evolutionary and epidemiological model predicts that under certain ART scenarios, which align closely with the “treatment as prevention,” “universal testing and treatment,” and “90-90-90” scenarios that HIV public health programs aspire to implement, HIV virulence may increase relatively rapidly. These results are seen both in a model calibrated based on South Africa HIV incidence and prevalence trends and in an alternate model with entirely different parameterization. We note that incidence declines in these scenarios of increased virulence, but individuals with untreated infections will progress more quickly and per-act transmission rates will rise. These results are consistent with theoretical explorations of the effects of treatment on pathogen virulence, a key observation of which was that increasing treatment rates resulted in increasing optimal virulence (33); we observed this same result, as increasing ART coverage resulted in increasing virulence. Our results are also qualitatively consistent with a recent study that used a deterministic model to predict the effects of ART on the relative frequencies of two HIV strains representing high or low virulence (34). Further study is warranted to assess these HIV-specific predictions, including modeling aspects of combination prevention programs that may modulate these effects, *e.g.* PrEP, medical male circumcision, or condom use, and using cohort data to evaluate empirical relationships between trends in ART coverage and HIV virulence markers.

An additional conclusion from our study is that standard HIV epidemic models (models that do not include parameters related to population-level variation in SPVL and to the capacity for HIV virulence to evolve) may underestimate the benefits of ART prevention programs. It may be beneficial, as HIV epidemiologic models continue to develop, to include realistic functions of viral evolutionary dynamics in such models. Even in worst case scenarios, our modeling study suggests that increasing virulence driven by universal test and treat is likely not a major public health concern, with a risk far outweighed by the benefits of improved clinical outcomes and reduced incidence. Changing virulence is amenable to being monitored alongside transmitted drug resistance in sentinel surveillance.

## MATERIALS AND METHODS

We previously developed a stochastic, agent-based HIV evolutionary and epidemic model that simulates viral dynamics within and between individuals (6). This model allows for an HIV virulence phenotype (set point viral load; SPVL) to change over the course of a simulated epidemic, and thus provides an evolutionary framework in which a balance can be achieved between infectivity (efficiency of viral transmission) and virulence (rate of disease progression). The underlying model was written in C with a front-end written in R. The code is freely available from the authors, upon request.

### Simulated population

Each epidemic simulation starts with *N* total individuals at time zero (HIV-uninfected and infected), with each infected individual (of *n* total HIV-infected individuals) provided a SPVL value randomly selected from a normal distribution with user-defined mean and variance (Table 1). Entry of new (HIV-uninfected) individuals into the population occurs at a constant rate, such that the overall population will stay at its initial value.

### Viral load parameters

The model includes the following parameters related to HIV viral load: 1) the distribution of SPVL in a population of HIV-infected individuals (13, 14); 2) the daily progression of viral load over the course of an individual infection, including distinct viral load trajectories for acute, chronic, and AIDS stages (15, 35-37); 3) the predictive relationship between HIV viral load and the per-day transmission rate (assuming a given probability of sexual contact per day; see below) (4, 5); 4) the predictive relationship between SPVL and the rate of disease progression, mediated through rates of individual CD4+ T cell decline that are stratified by individual SPVL (38); and 5) a viral role in the determination of each individual’s SPVL (*i.e.* variation in the viral genotype explains a portion of the population variation in SPVL; non-zero heritability of SPVL in the infected population) (20-24). Parameter estimates for these viral components were fixed based on relevant literature (Table 1), and parameter uncertainty was addressed systematically by sensitivity analyses reported in our previous description of the model (6), and by performing our experiments on two separate model calibrations. See Supporting Information for further descriptions of the above model functions.

### Primary model calibration

We performed a two-step process to calibrate our primary model, with the intent to reproduce incidence and prevalence trajectories based on prevalence data from South Africa (39), so that the output of our main analysis was directly comparable to the 12 HIV epidemic models described in Eaton *et al.* (27). As such, we first used evidence-based (*i.e.* viral load, CD4) parameter values within our evolutionary model; these are discussed in depth in our previous description of the model, and in the Supporting Information (6). Second, we calibrated assumption-based (*i.e.* behavioral) parameter values specifically to produce epidemic trends similar to those observed in South Africa, and based on similar calibration of the 12 models included in Eaton *et al.* (27).

It is not straightforward to calibrate HIV epidemic models to accurately reflect the decreases in incidence and prevalence observed in epidemics of sub-Saharan Africa (40); as in all epidemic models, it is widely accepted that the observed epidemic trends in sub-Saharan Africa require relatively complex assumptions about population structure (different risk groups), patterns of sexual contact, or changes in risk behavior over time (40, 41). The 12 models in Eaton *et al.* all dealt with this issue, and used varying degrees of assumption-based parameter values in their calibration; none were able to reproduce realistic epidemic dynamics without some underlying epidemiological complexity (27). Our choice of behavioral parameters to include in this primary model, based on an epidemic with a core group of individuals with increased transmission rates, and the parameter settings of which produced the calibrated model output, follow from previous HIV epidemic models. This approach allowed us to externally validate our model epidemic output, and to interpret potential clinical and epidemiological impacts of our evolutionary output in a realistic (and accepted) framework. Further explanation of our model parameterization and calibration is included in the Supporting Information.

### Alternate model calibration

It was not the goal of our overall study to assess the potential effects of variation in behavioral parameters on the interaction of ART and HIV virulence evolution. Rather, our goal was to asses the effects of ART on HIV virulence evolution, with ART applied under a wide variation of scenarios and coverage, while maintaining the primary epidemic calibration based on South Africa and the 12 models in Eaton *et al.* However, we assessed whether the predicted effects of ART on virulence evolution were robust to variation in the overall epidemic scenarios (different variants of the underlying epidemic model, including underlying parameterization and resulting incidence and prevalence trends). To do this, we modified our behavioral parameters by: 1) decreasing the mean degree; 2) removing the core group of individuals that had short relative relationship durations and elevated rates of sexual contact; 3) incorporating a random mixing sexual network that eliminated the assortative mixing of sexual contacts by relationship duration category (Table 1). With this alternate epidemic model we repeated the entirety of our ART-based experiments.

### Antiretroviral therapy parameters

We structured ART dynamics as a heuristic starting point for theoretical studies of ART and HIV virulence evolution. As such, we assumed a single, standard regimen, with complete adherence and retention, and without the emergence of drug resistant mutations and associated changes in viral fitness (numerous studies report that transmitted drug resistance is rare in most populations (42, 43) and that the vast majority of transmitted drug resistance mutations have low fitness costs (44, 45). Individual ART use was applied after fulfilling necessary criteria of *eligibility threshold*, *initiation time*, and *population coverage*. We evaluated two types of ART eligibility scenarios. First, eligibility was based on CD4 count thresholds (entry into a CD4 count category: CD4>500; CD4<500; CD4<350; CD4<200). These scenarios are most relevant to how treatment has been implemented in recent years, as countries have followed WHO guidelines. Second, eligibility was based on time elapsed since date of infection (one, two, three, four, five, or six years after infection). These scenarios reflect likely changes in the future as treatment becomes nearly universally available through the UNAIDS 90-90-90 initiative.

Initiation of ART after an individual became eligible was immediate for the “time since infection” eligibility criterion but was delayed for one year for the “CD4 count category” criterion (this is consistent with the models compared in Eaton *et al.* (27), and realistic given testing rates). No individuals were eligible for ART during acute infection in any scenario (regardless of CD4 count category). Population-level ART coverage was implemented via individual probabilities of initiating ART, after meeting the eligibility criteria described above. (With complete adherence, our estimates of population-level coverage are likely slight overestimates of coverage rates in epidemics with equivalent individual probabilities of initiating ART but less than complete adherence.) Decreases in transmission probability for individuals receiving ART were mediated entirely by a immediate decrease in viral load to 50 copies/mL upon treatment initiation.

### Simulations and output

We ran epidemic simulations for 60 years, in discrete time-steps of one day. For each model run we tracked the distribution (mean, median and variance) of SPVL (viral load at the end of acute infection and prior to initiation of ART) and the population incidence and prevalence. For specific comparisons to the models described in Eaton *et al.* (27), we evaluated changes to mean SPVL, incidence, and person-years of ART per infection averted that were observed at eight and 38 years after the roll-out of a population-level ART program that began in 2012—with treatment eligibility at CD4 count <350 cells/uL and population coverage at 80% (compared to counterfactuals in the same populations without ART). This specific treatment scenario was used by Eaton *et al.* (27) to approximate an implementation of World Health Organization guidelines (current at the time of that study) and the Joint United Nations Programme on HIV/AIDS definition of ‘‘universal access’’ as reaching 80% of HIV-infected individuals.

Thus, we began our simulations in year 1990, ART was introduced to the populations in the beginning of year 2012, and evaluations were done using output from the midpoint of years 2020 (eight years) and 2050 (38 years). In addition to the specific comparison to Eaton *et al.* outputs, we evaluated combinations of ART coverage (40 to 100%, by 20% increments) and either ART time since infection eligibility (one, two, three, four, five, or six years after infection) or ART CD4 count threshold eligibility (all eligible (CD4>500), CD4<500, CD4<350, or CD4<200). For each combination of coverage and eligibility we performed 10 replicate simulations.

## ACKNOWLEDGMENTS

This work was supported by grants from the U.S. National Institutes of Health (R01AI108490 to J.T.H., J.E.M., and S.G., and P30AI027757 to the University of Washington Center for AIDS Research). The content is solely the responsibility of the authors and does not necessarily represent the official views of the National Institutes of Health. The funding sources had no role in the writing of this manuscript or the decision to submit it for publication.

## Supporting Information

### Additional details of the epidemic model

#### Within-host component (viral load dynamics)

For each individual, upon HIV-1 infection with an initial viral population of size *V*_0_, viral load increases exponentially at rate *r* until a user-defined peak viremia, *V*_peak_, at a user-defined time *t*_peak_. Viral load then declines exponentially at an individual-specific decay rate, *d*_acute_, until it reaches set point viral load (SPVL), *V*_sp_, at time *t*_sp_. We define *t*_sp_, the time when *V*_sp_ is established, in the number of days after infection; this is a user-defined length from initial infection through peak viremia until set point is reached. After reaching set point (*t*_sp_), the viral load increases as (Equation 1):

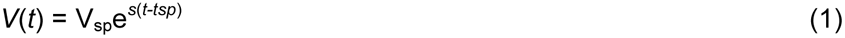

where *s* is the user-defined parameter of annual viral load increase rate, and *t* is the time that has elapsed since the patient was first infected. For all HIV-1 infected individuals, we assume a log linear change in viral load after SPVL has been established until the onset of AIDS. Functionally this can be set to zero change, equivalent to stable viral load in the asymptomatic stage. Viral load upon onset of AIDS is defined as the same for all individuals, and is independent of SPVL. To reconcile population variation in SPVL (*V*_sp_) with primary infection dynamics, we assume that *V*_0_, *r*, *t*_peak_, *t*_sp_ and *s* are the same in all individuals, but that *d*_acute_ = *ln*(*V*_peak_/*V*_sp_)/(*t*_sp_ - *t*_peak_) varies.

#### Within-host parameters for disease progression (CD4+ T-cell count parameters)

To model the relationship between SPVL and the rate of disease progression, as well as incorporate ART initiation thresholds based on CD4+ T cell counts, we included an intermediary function that related individual SPVL to the starting CD4 count category (individual CD4 count immediately after infection), and to subsequent disease progression based on waiting times in four CD4 count categories: CD4>500; 500>CD4>350; 350>CD4>200; CD4<200 (AIDS). This function is based on data from the AIDS Therapy Evaluation in the Netherlands (ATHENA) observational cohort, which includes HIV-infected individuals followed in the 27 HIV treatment centers in the Netherlands since 1996 [27].

#### Across-host component (viral load and transmission)

We assume that transmission rates follow available data from serodiscordant heterosexual partners; the probability of a HIV-infected person will transmit to a HIV-negative person is determined by an increasing Hill function (Equation 2) that follows Fraser [4]:

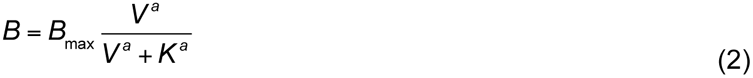

where *B*_max_ is the maximum probability of transmission per year, *V* is the donor’s viral load at the time of sexual contact, *a* is a Hill Coefficient that influences the steepness of the response curve, and *K* is the viral load at which the probability of transmission to a susceptible person is *B*_max_/2. We have followed the parameter values from Fraser [4], except for *B*_max_, which we have increased (from 0.001 per day in Fraser [4]) to account for the slower epidemic growth rates than expected, which may be explained as this value being from serodiscordant couples that are enrolled in HIV-1 cohorts and likely an underestimate relative to the general population.

#### Across-host component (heritability of set point viral load)

The SPVL, *V*_sp_, is determined by both viral (the viral genotype, *VirCont*) and environmental (a combination of undefined host and environmental factors, *EnvCont*) factors. *VirCont* is different for each individual; HIV-infected individuals inherit the value of *VirCont* from the individual who infected them, with random variance introduced within a separate mutational variance parameter. Mathematically, we assume, for donor (*i*) and recipient (*j*) (Equations 3 - 5):

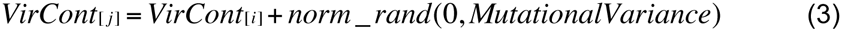

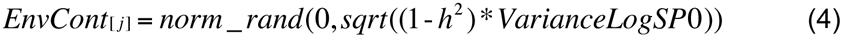

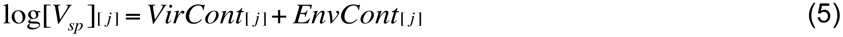

where *h*^2^, the user-defined heritability.

#### Sexual mixing network

For each individual in a simulation, the model maintains a list of sexual partnerships and viral transmission pairs. Individuals were not marked by sex, and risks of infection were bidirectional, meaning that both partners have independent probabilities of infecting the other partner that are dependent on individual viral loads.

For any one simulation, prior to the simulation of viral transmission, an initial set of contacts is formed by randomly choosing pairs from the population. The probability of each person entering into this link is set to *e*^-Li^, where *L*_i_ is the number contacts that person *i* has (for this analysis we did not allow for concurrent relationships). If this probability was not met (or if *L*_i_ > *MaxLinks*), another partner is selected at random from the population. This process is repeated until the total number of links equals *N*\**M*/2, where *N* is the total number of sexual active individuals, and *M* is the mean degree. The probability of a connection between individuals *i* and *j* dissolving is set to 2/(*Duration*[*i*] * *Duration*[*j*]), where *Duration*[*x*] is the expected time that person *x* stays in a relationship. After removing links from all newly dissolved partnerships, *N*\**M*/2 - *L*/2 links are added to the system, where *N* is the number of individuals after accounting for births and deaths that occurred that day and *L* is the number of links in the system after the dissolution step. If this quantity is negative, no links are added.

#### Behavioral parameters

The virologic and CD4-based progression functions were embedded in a host population without explicit demographic (sex, age) heterogeneity, but with behavioral heterogeneity. Each individual was assigned a relational duration propensity (range: 6-60 months); when two individuals partner, their relationship was slated to last for the mean of these individual effects. The daily probability of sex varied by relationship duration (100% for relationships <= 6 months; 5% for relationships >6 months and <=30 months; 3% for relationships >30 months). The coital frequencies for the two higher categories were selected as part of the process of calibrating the model to epidemic data. Individuals were placed into relationship partnerships randomly, with no concurrency (those already in a relationship were not eligible to form a new partnership). The rate of partnership formation was a calibrated parameter value. We did not include a change in behavior over time, *e.g.* a reduction in the sexual contact rate over calendar time.

**Supporting Information Figure S1.**
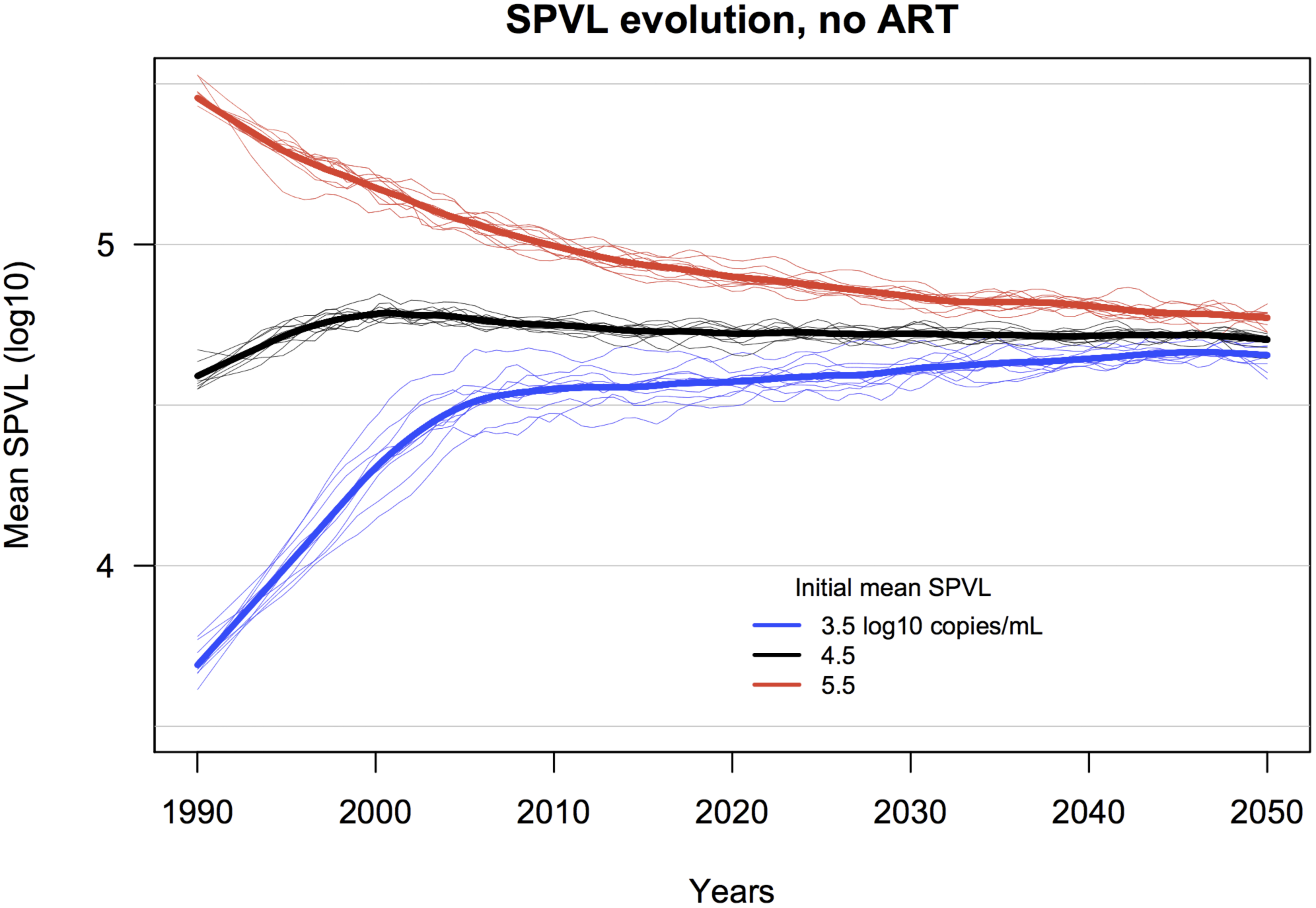
Simulated trends in HIV virulence (via set point viral load (SPVL) as a proxy) reveal adaptive evolution toward an optimum SPVL. Shown are LOESS regression lines for ten random replicates for each initial mean SPVL (thin lines), and the mean of these replicates (thick lines).

**Supporting Information Figure S2.**
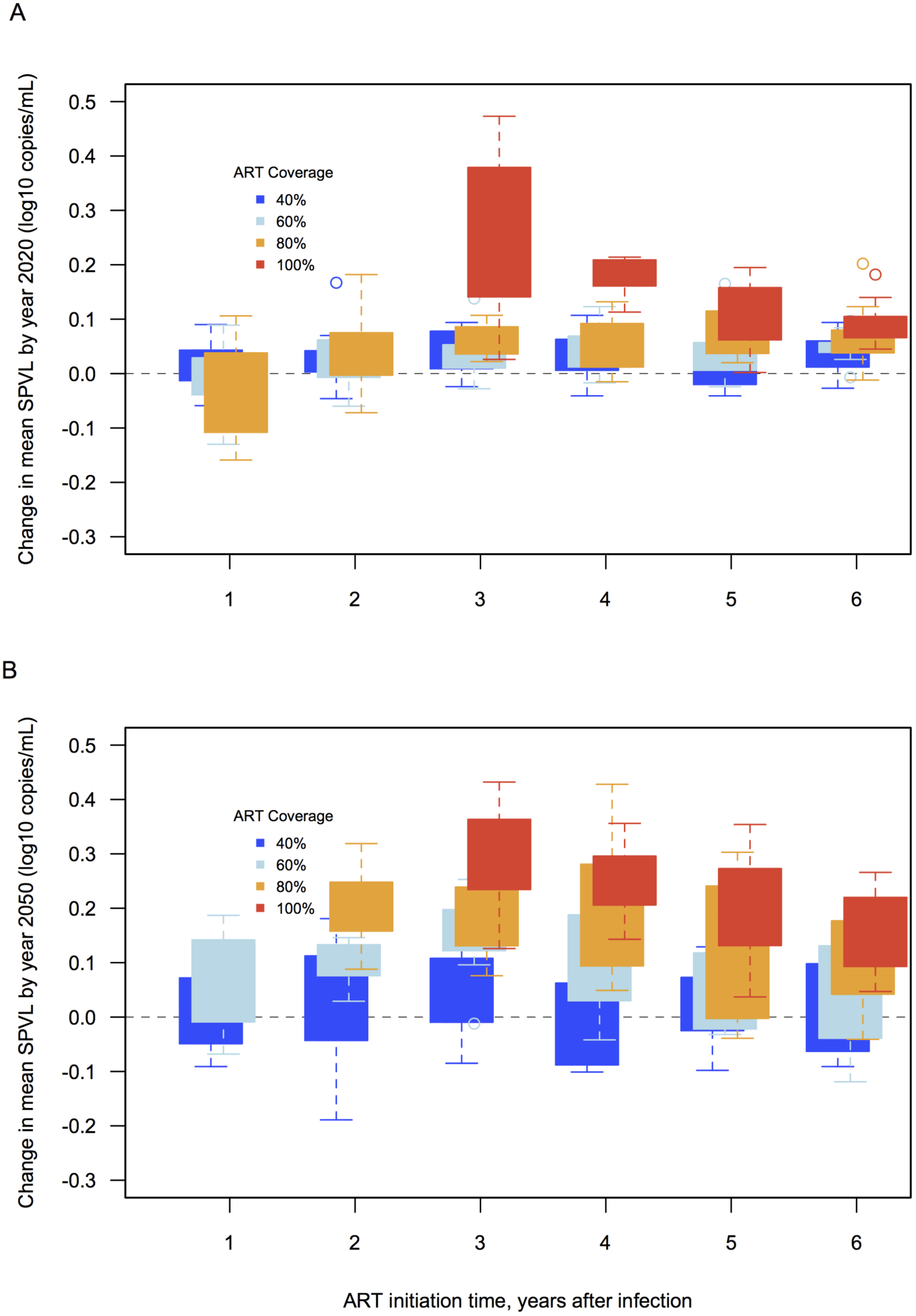
Simulated effects of ART on HIV virulence evolution, measured at **A)** 8 and **B)** 38 years after rollout of ART (*i.e.* from ART rollout in 2012 to 2020 and 2050). ART initiation is determined based on time elapsed since infection. (Increases in mean SPVL for 100% coverage are not reported for initiation at one and two years after infection because incidence is 0 in those scenarios.)

**Supporting Information Figure S3.**
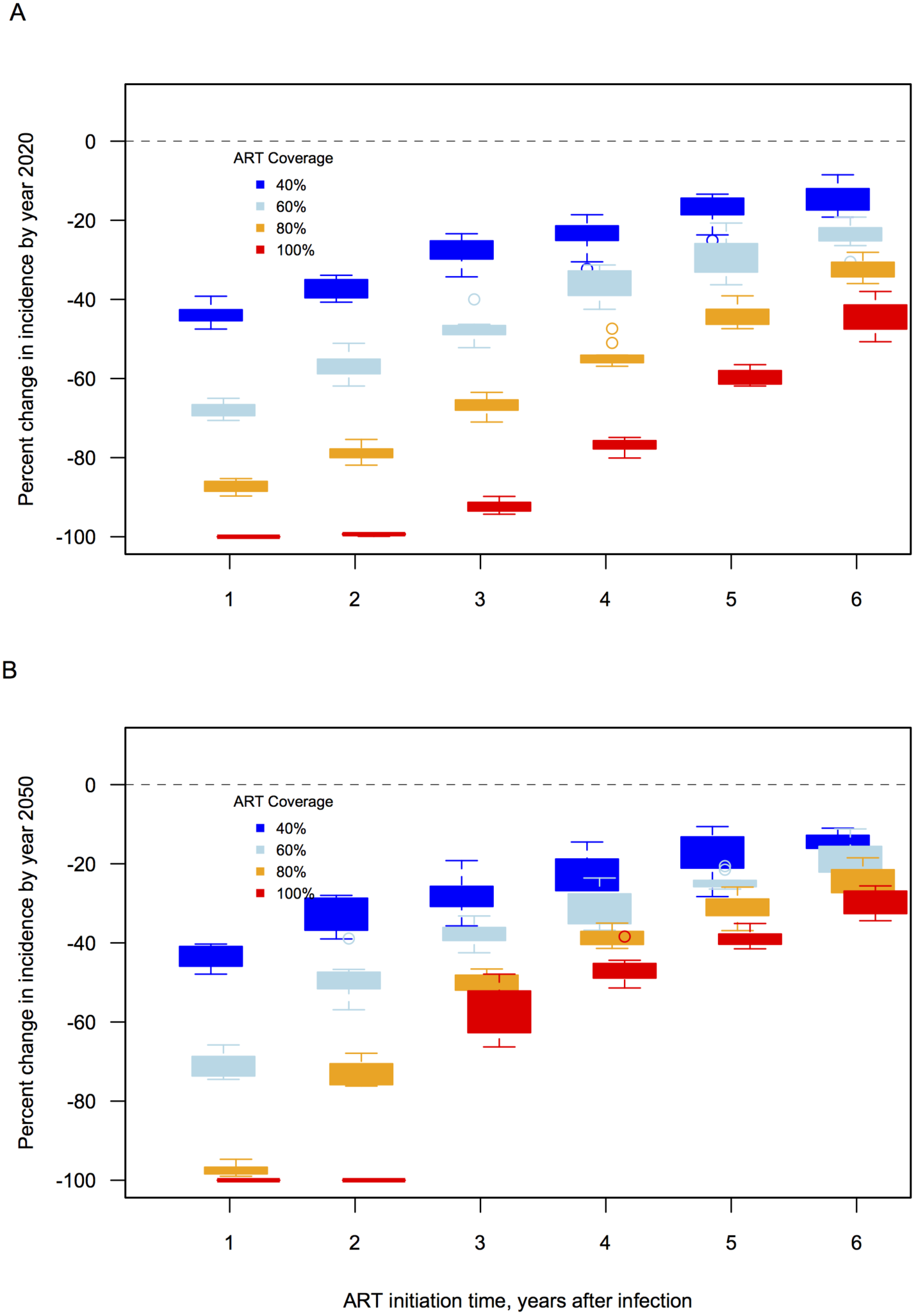
Simulated effects of ART on HIV incidence, for scenarios when ART is initiated based on time since infection, at increasing coverage levels, measured at **A**) Eight and **B)** 38 years after rollout of ART (from 2012 to 2020 and 2050). ART initiation is determined based on time since infection.

**Supporting Information Figure S4.**
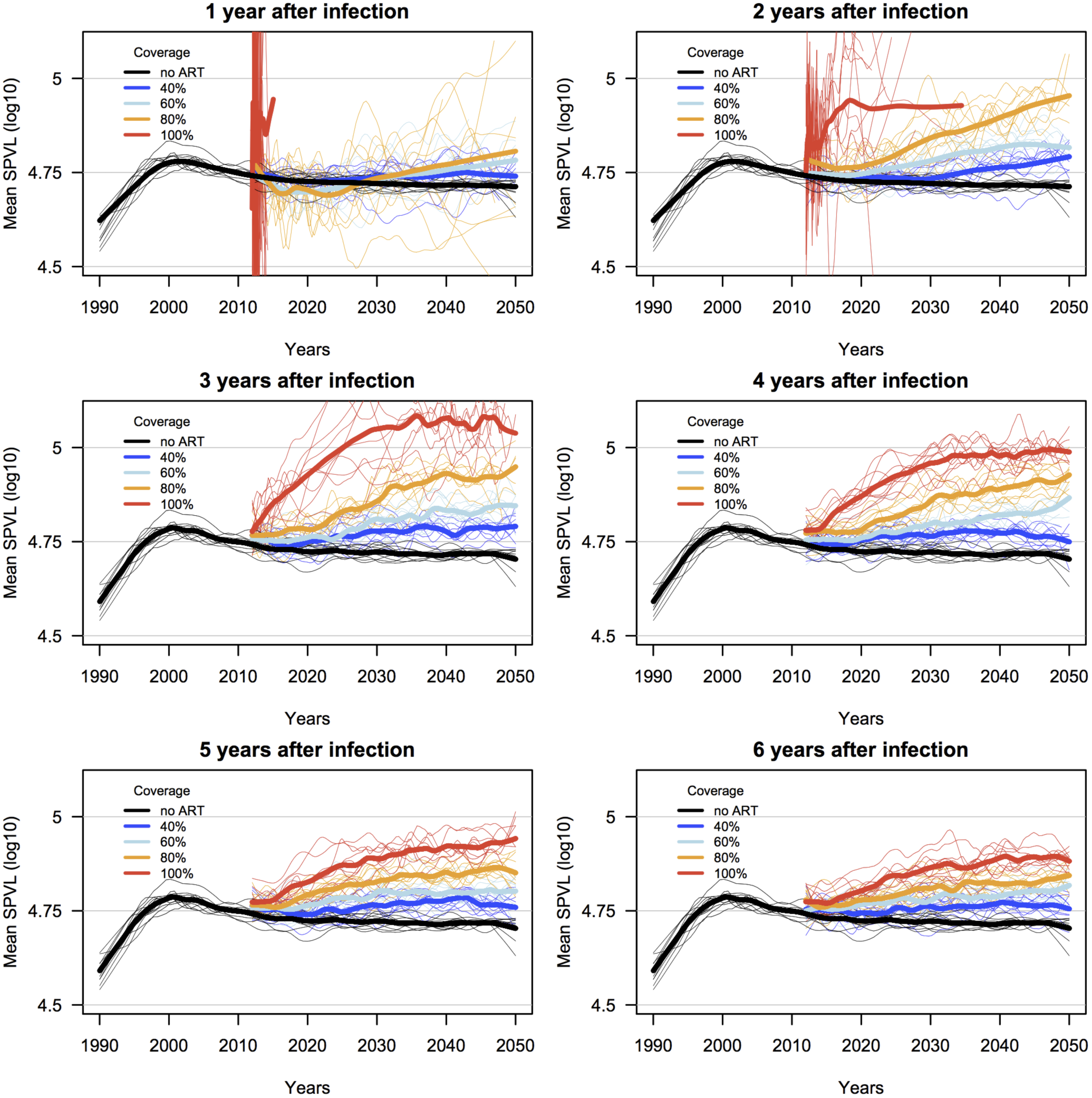
Simulated effects of ART on HIV virulence evolution, for scenarios when ART is initiated based on time since infection, at increasing coverage levels. Shown are LOESS regression lines for ten random replicates for each ART coverage scenario (thin lines), and the mean of these replicates (thick lines). Initial mean SPVL was 4.5 log_10_ copies/mL. ART coverage lines start at year 22, corresponding to a simulation starting at year 1990 with ART rollout at year 2012. Truncated lines are the result of epidemic runs ending with 0% incidence.

**Supporting Information Figure S5.**
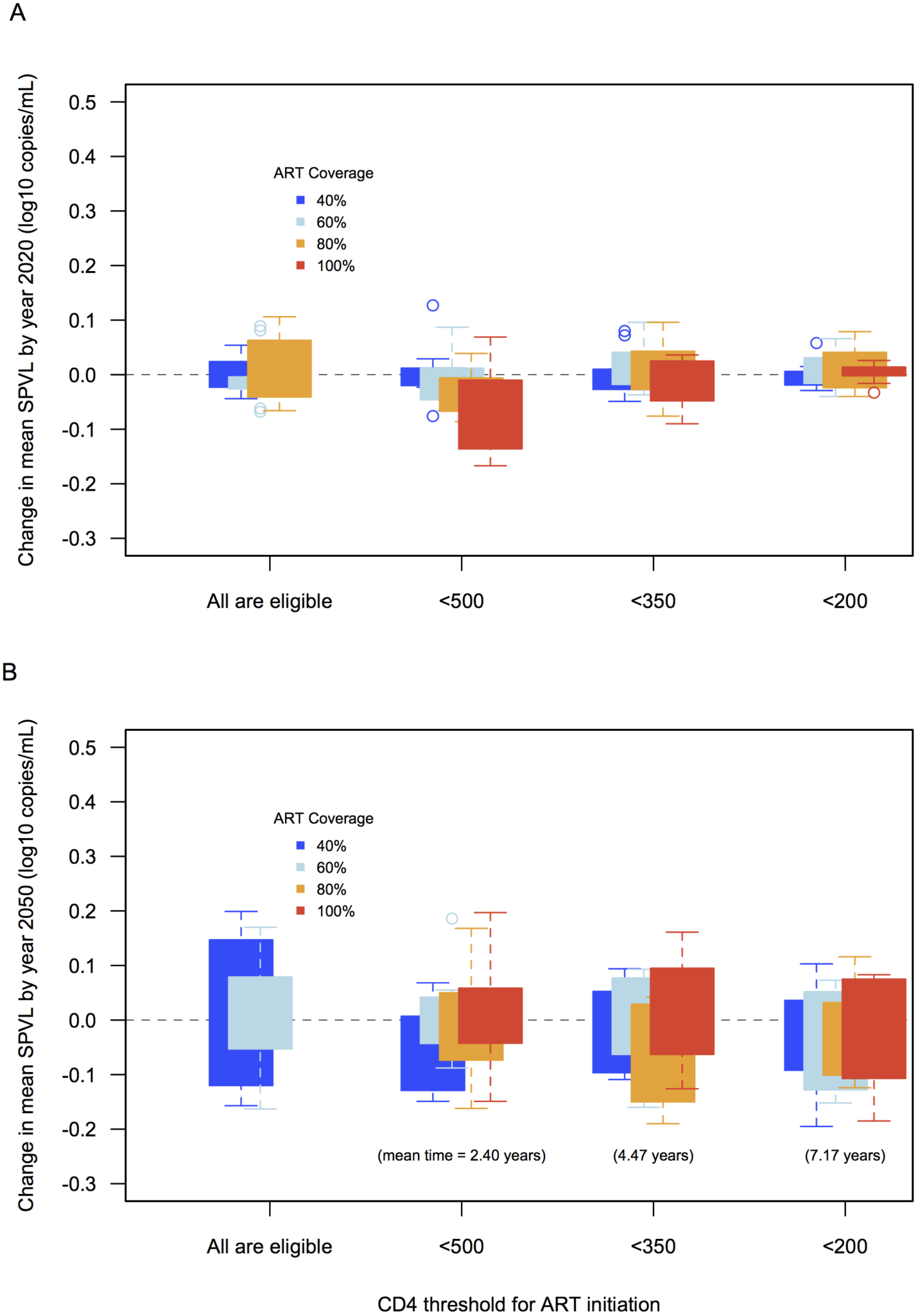
Simulated effects of ART on HIV virulence evolution, measured at **A)** 8 and **B)** 38 years after rollout of ART (from ART rollout in 2012 to 2020 and 2050). ART initiation is determined based on CD4 count eligibility thresholds (shown on the X-axis). (Increases in mean SPVL for 100% coverage are not reported for CD4>500 because incidence is 0% in those scenarios.)

**Supporting Information Figure S6.**
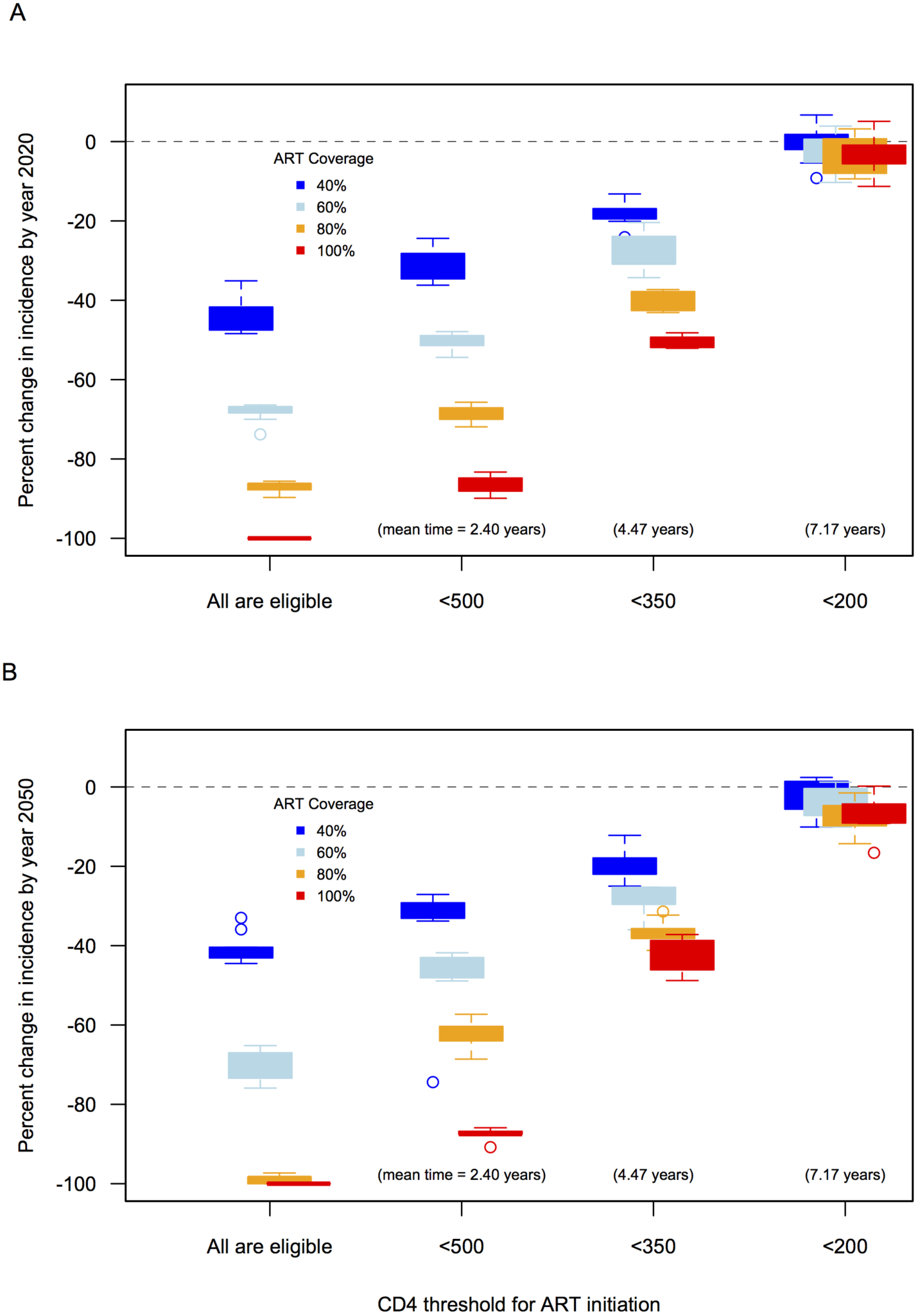
Effects of ART on HIV incidence, measured at A) eight and B) 38 years after rollout of ART (from 2012 to 2020 and 2050). ART initiation is determined based on CD4 count eligibility thresholds.

**Supporting Information Figure S7.**
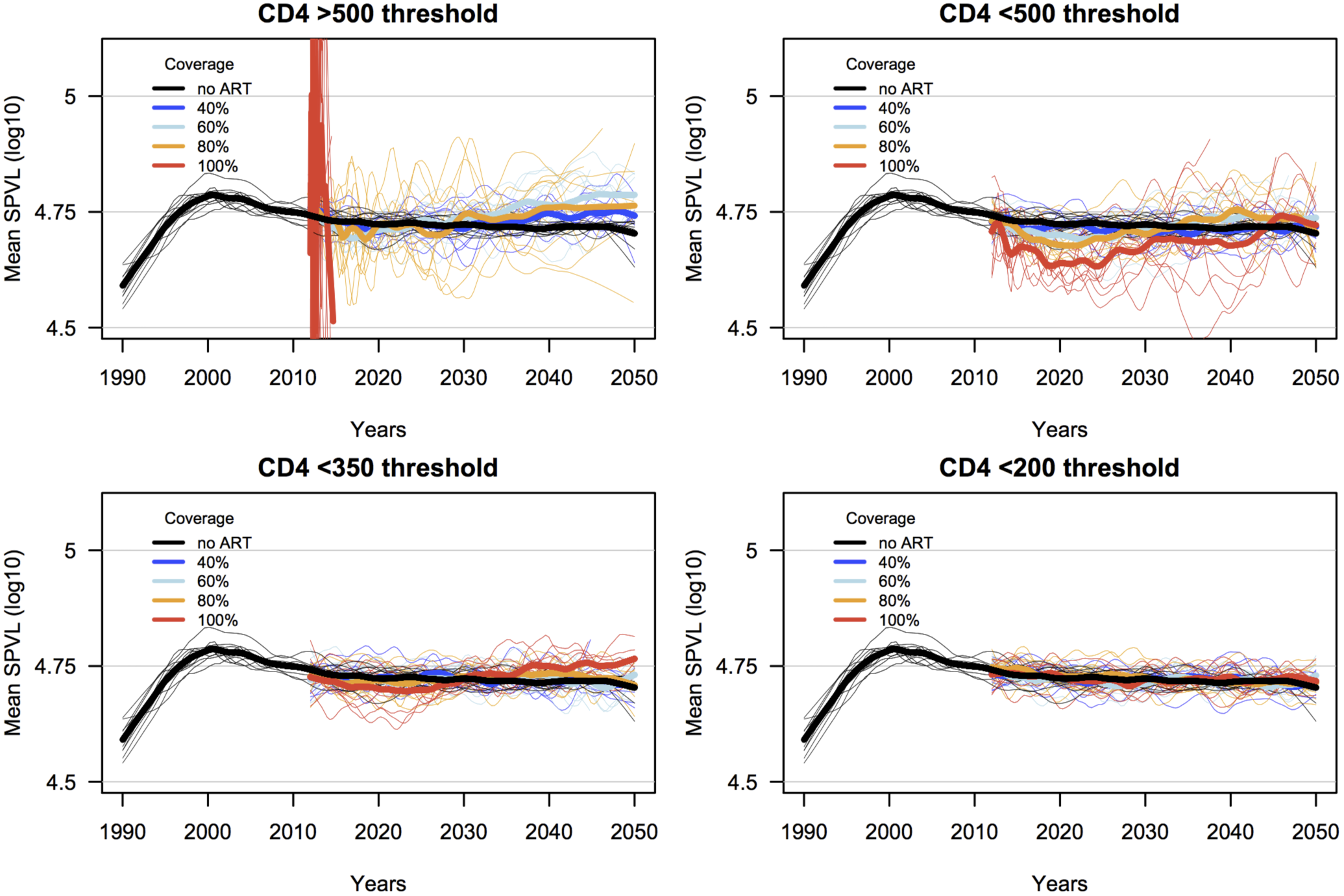
Simulated effects of ART on HIV virulence evolution, for scenarios when ART is initiated based on CD4+ T cell count, at increasing coverage levels. Shown are LOESS regression lines for ten random replicates for each ART coverage scenario (thin lines), and the mean of these replicates (thick lines). Initial mean SPVL was 4.5 log_10_ copies/mL. ART coverage lines start at year 22, corresponding to a simulation starting at year 1990 with ART rollout at year 2012. Truncated lines are the result of epidemic runs ending with 0 incidence.

**Supporting Figure S8.**
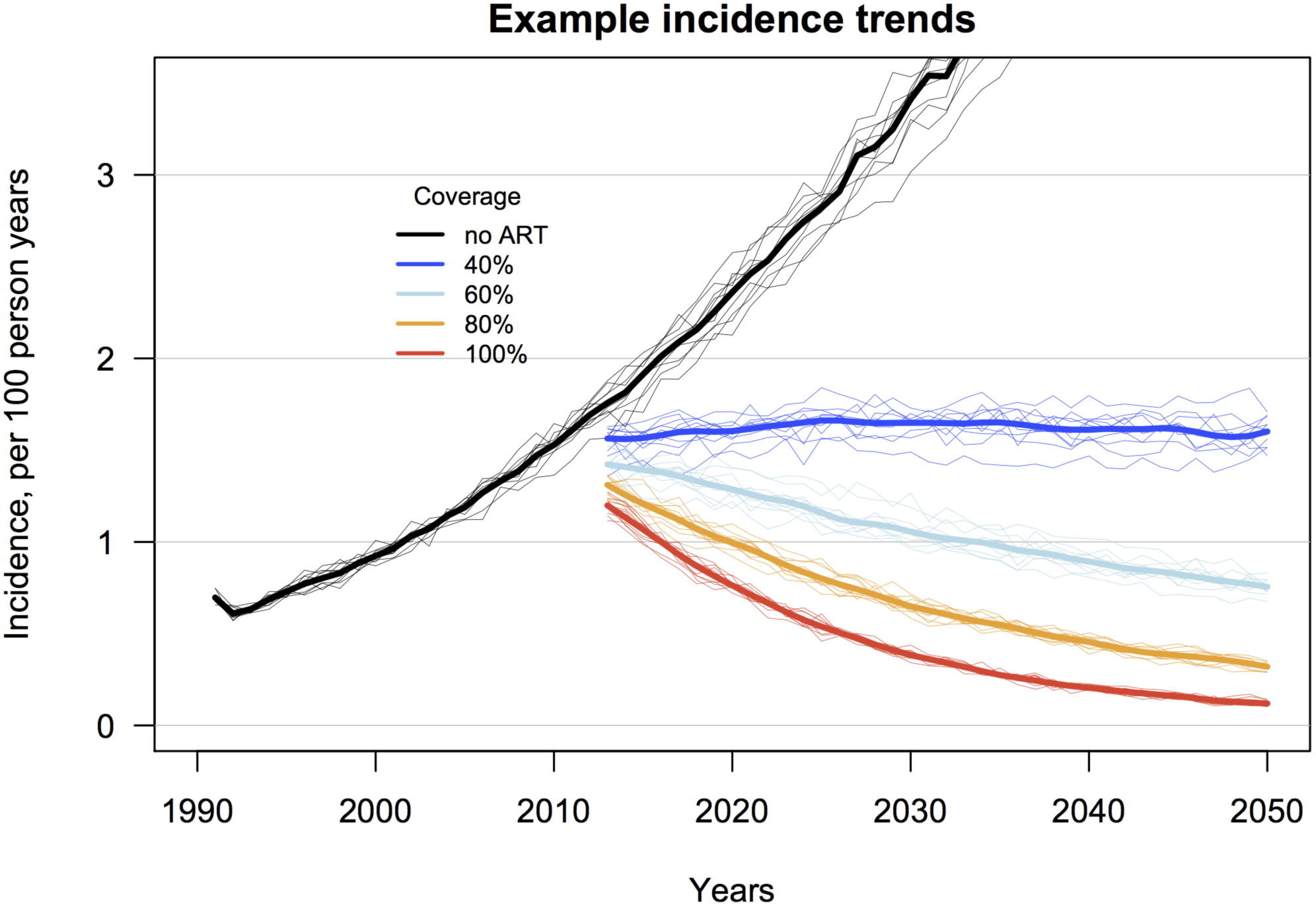
Alternate model results: simulated trends in HIV incidence for ART scenarios of 40, 60, 80 and 100% coverage (individual probability of receiving ART with complete adherence) and CD4 count threshold for treatment initiation <350 cells/mL, versus the counterfactual epidemic simulation with no ART. Shown are LOESS regression lines for ten random replicates for each ART coverage scenario (thin lines), and the mean of these replicates (thick lines). Initial mean SPVL was 4.5 log_10_ copies/mL.

**Supporting Information Figure S9.**
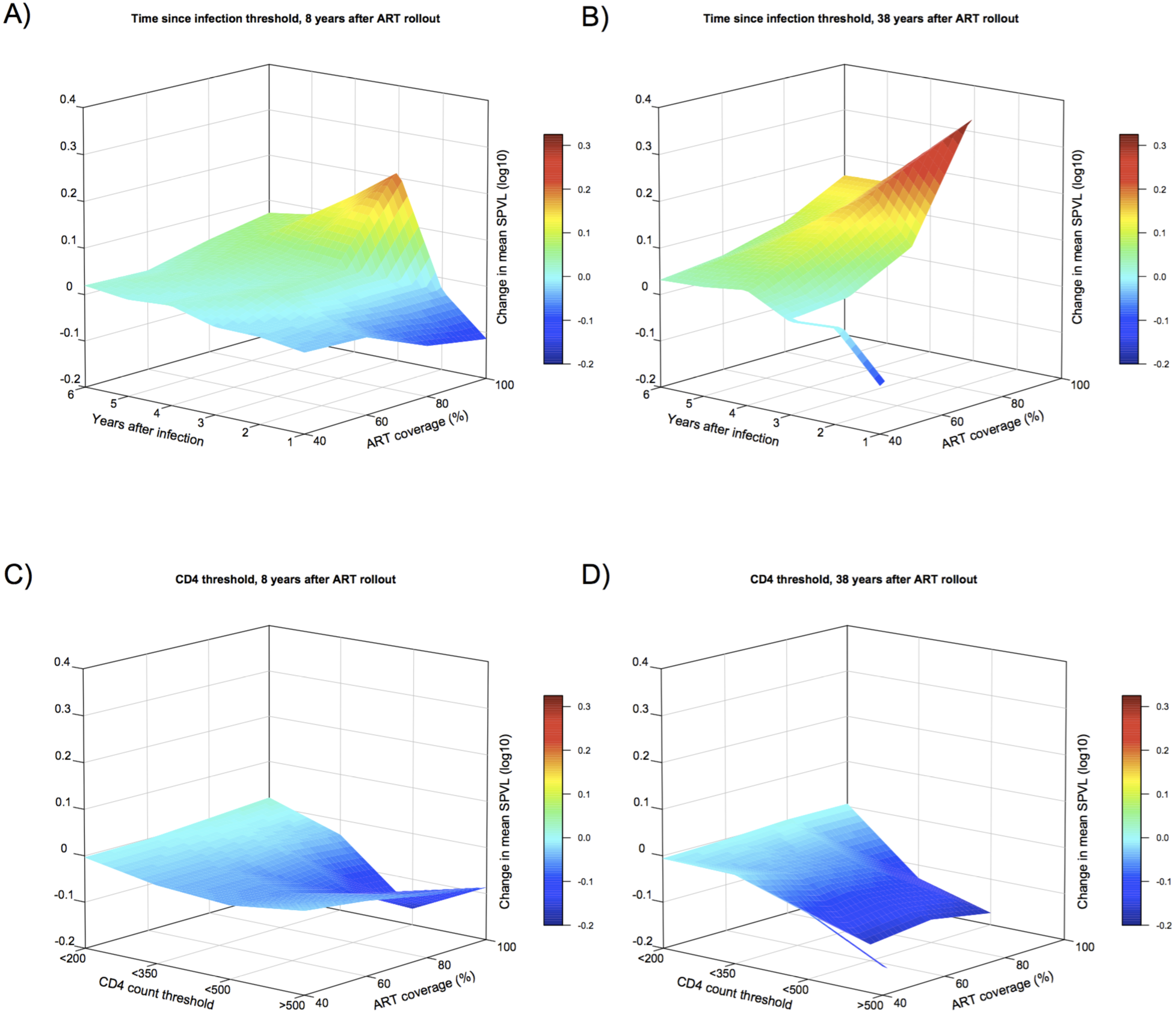
Alternate (different calibration) model results: surface plots showing change in mean SPVL between epidemic simulations with and without ART, for scenarios of increasing ART coverage (individual treatment probability) and ART initiation based either on time since infection or CD4 count threshold. For epidemic scenarios with ART initiation based on time since infection, **A)** shows mean SPVL change 8 years after ART rollout (from year 2012 to year 2020), and **B)** shows mean SPVL change 38 years after rollout (from year 2012 to year 2050). For epidemic scenarios with ART initiation based on CD4 count, **C)** and **D)** show mean SPVL at 8 and 38 years after rollout, respectively.

**Supporting Information Figure S10.**
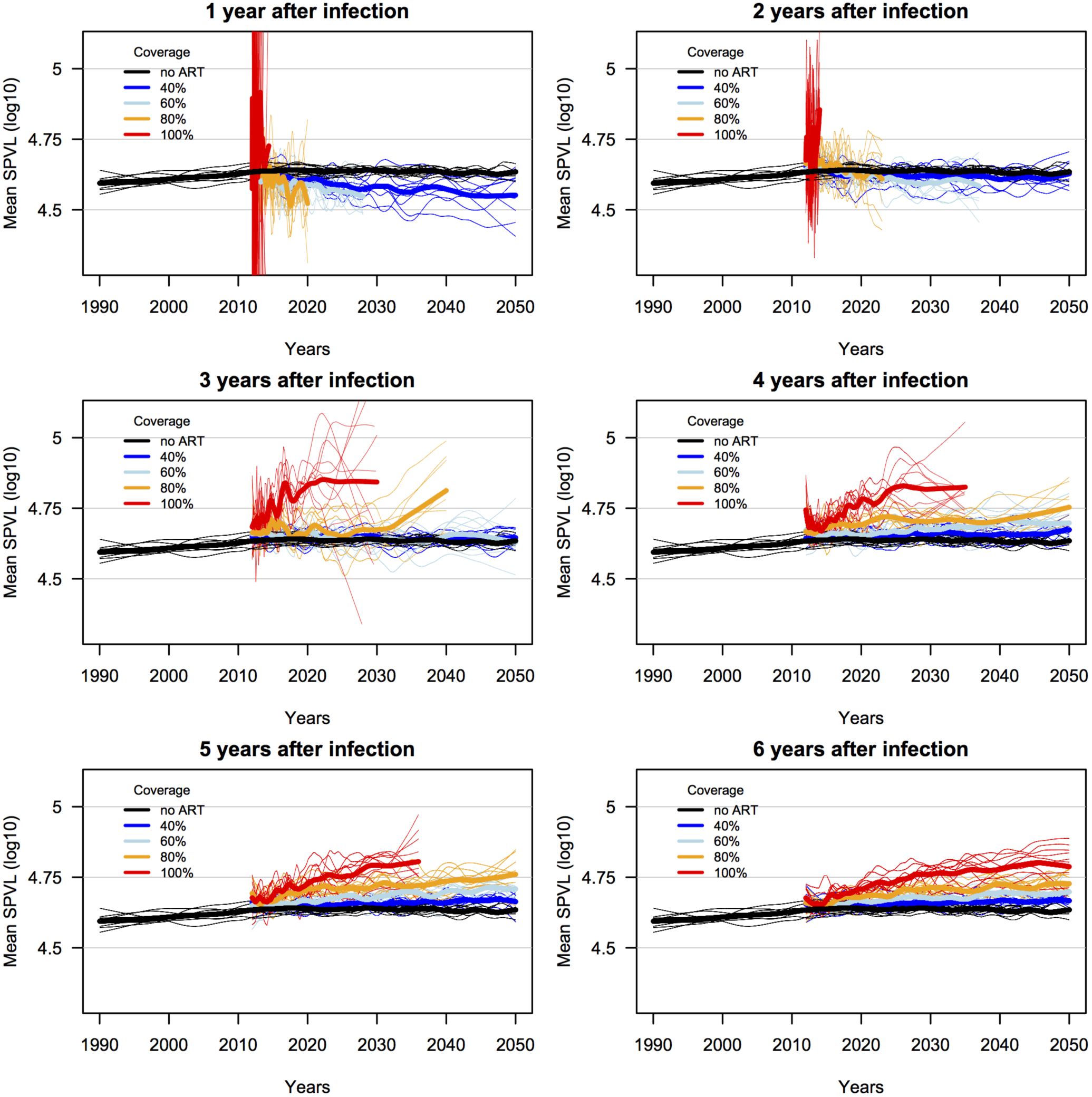
Alternate model results: simulated effects of ART on HIV virulence evolution, for scenarios when ART is initiated based on time since infection, at increasing coverage levels. Shown are LOESS regression lines for ten random replicates for each ART coverage scenario (thin lines), and the mean of these replicates (thick lines). Initial mean SPVL was 4.5 log_10_ copies/mL. ART coverage lines start at year 22, corresponding to a simulation starting at year 1990 with ART rollout at year 2012. Truncated lines are the result of epidemic runs ending with 0 incidence.

**Supporting Information Figure S11.**
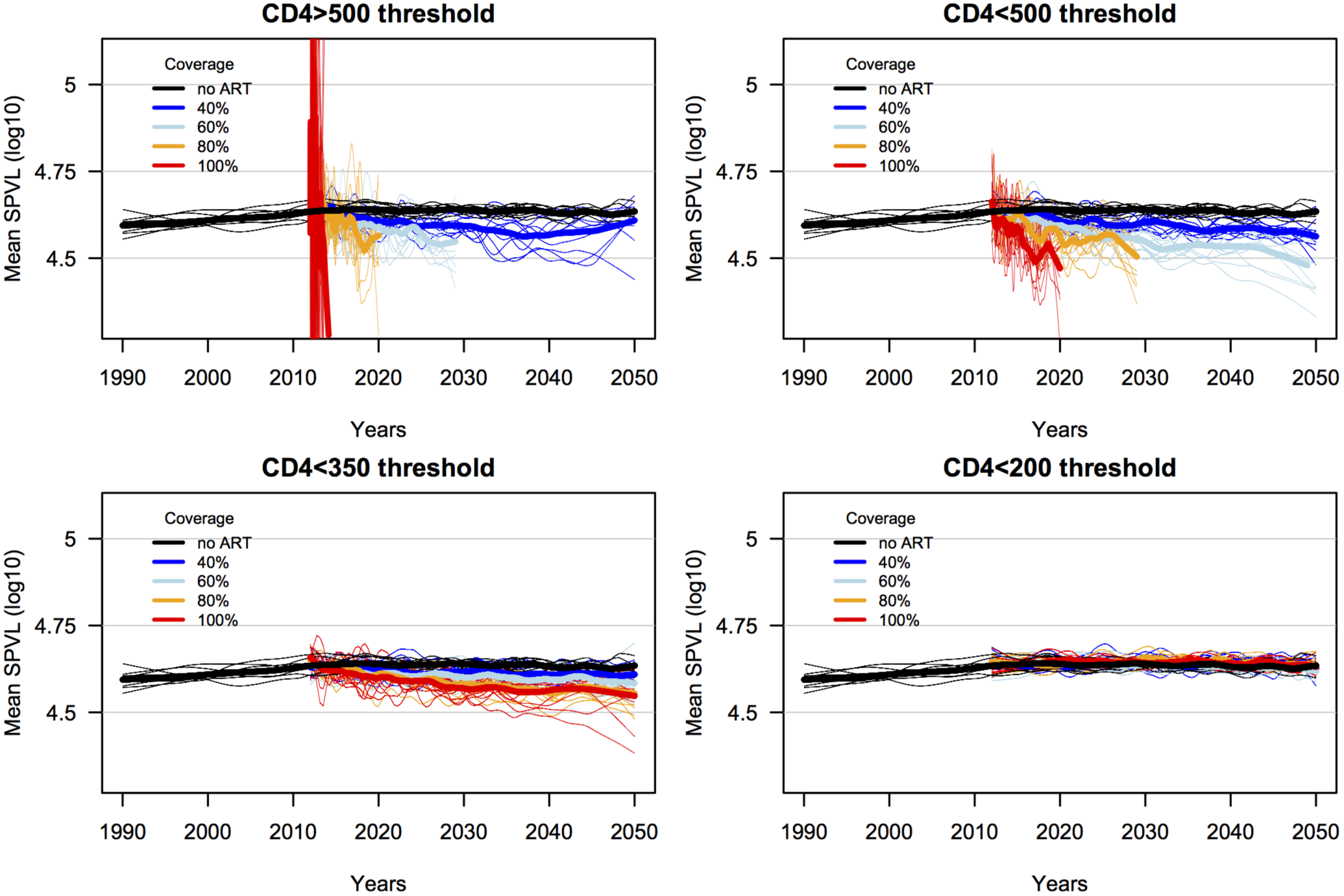
Alternate model results: simulated effects of ART on HIV virulence evolution, for scenarios when ART is initiated based on CD4+ T cell count, at increasing coverage levels. Shown are LOESS regression lines for ten random replicates for each ART coverage scenario (thin lines), and the mean of these replicates (thick lines). Initial mean SPVL was 4.5 log_10_ copies/mL. ART coverage lines start at year 22, corresponding to a simulation starting at year 1990 with ART rollout at year 2012. Truncated lines are the result of epidemic runs ending with 0 incidence.

**Supporting Information Figure S12.**
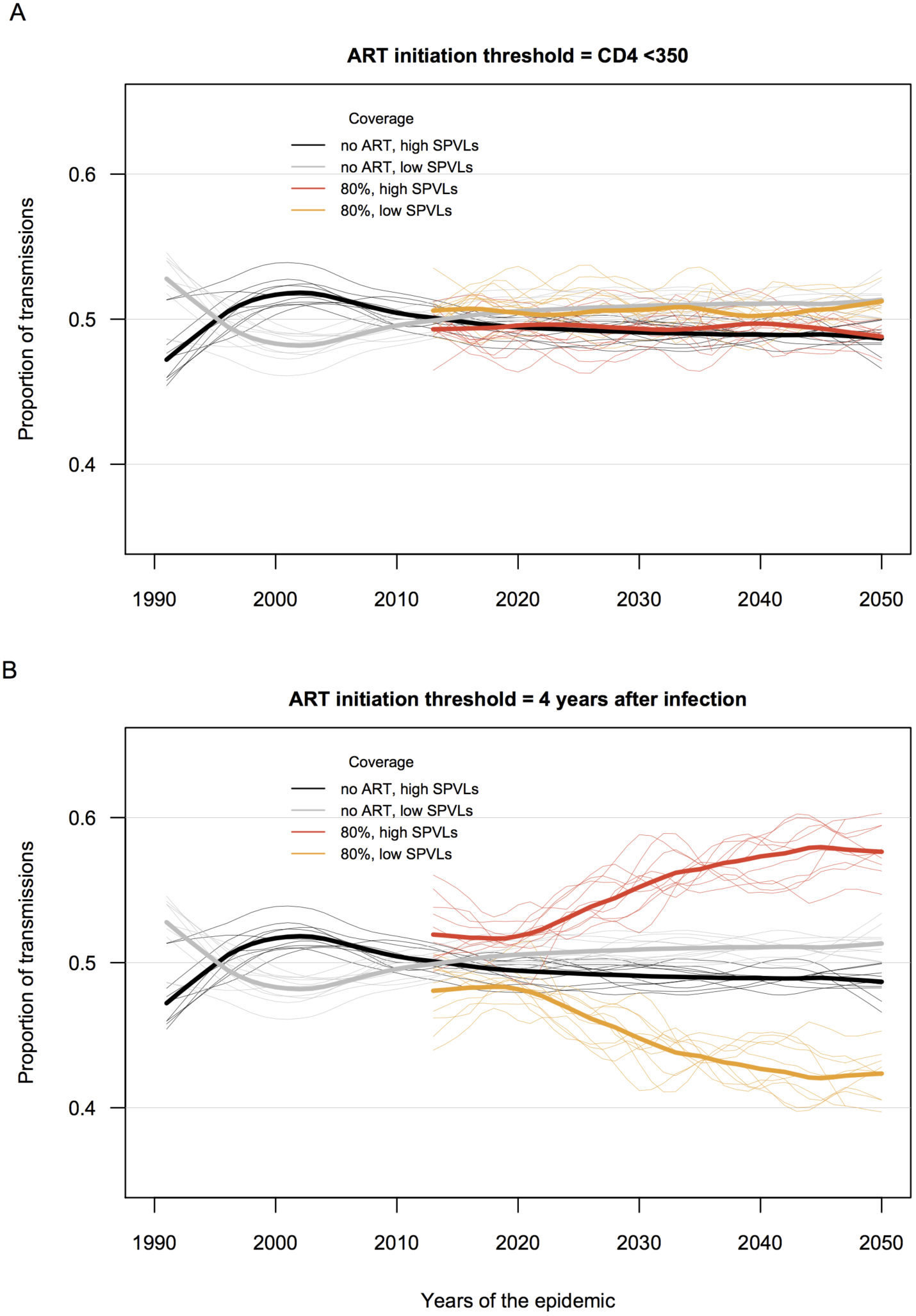
Proportion of transmissions from SPVL lineages with higher (>4.74 log_10_ copies/mL) and lower (<4.74 log_10_) levels than the population mean prior to ART rollout, for a scenario of 80% ART coverage and two different ART initiation scenarios: A) ART eligibility based on CD4<350; or B) ART eligibility at four years elapsed after infection.

## REFERENCES

1. UNAIDS. The Gap Report: UN Joint Programme on HIV/AIDS UNAIDS. 2014.

2. Lundgren JD, Babiker AG, Gordin F, Emery S, Grund B, Sharma S, et al. Initiation of Antiretroviral Therapy in Early Asymptomatic HIV Infection. N Engl J Med. 2015;373(9):795–807.

3. Cohen MS, Chen YQ, McCauley M, Gamble T, Hosseinipour MC, Kumarasamy N, et al. Prevention of HIV-1 infection with early antiretroviral therapy. N Engl J Med. 2011;365(6):493–505.

4. Fraser C, Hollingsworth TD, Chapman R, de Wolf F, Hanage WP. Variation in HIV-1 set-point viral load: epidemiological analysis and an evolutionary hypothesis. Proc Natl Acad Sci USA. 2007;104(44):17441–6.

5. Shirreff G, Pellis L, Laeyendecker O, Fraser C. Transmission selects for HIV-1 strains of intermediate virulence: a modelling approach. PLoS Comput Biol. 2011;7(10):e1002185.

6. Herbeck JT, Mittler JE, Gottlieb GS, Mullins JI. An HIV epidemic model based on viral load dynamics: value in assessing empirical trends in HIV virulence and community viral load. PLoS Comput Biol. 2014;10(6):e1003673.

7. Anderson RM, May RM. Coevolution of hosts and parasites. Parasitology. 1982;85 (Pt 2):411–26.

8. Ewald P. Host-parasite relations, vectors, and the evolution of disease severity. Ann Rev Ecol Syst. 1983;14:465–85.

9. de Wolf F, Spijkerman I, Schellekens PT, Langendam M, Kuiken C, Bakker M, et al. AIDS prognosis based on HIV-1 RNA, CD4+ T-cell count and function: markers with reciprocal predictive value over time after seroconversion. AIDS. 1997;11(15):1799–806.

10. Mellors JW, Kingsley LA, Rinaldo CR, Jr., Todd JA, Hoo BS, Kokka RP, et al. Quantitation of HIV-1 RNA in plasma predicts outcome after seroconversion. Ann Intern Med. 1995;122(8):573–9.

11. Mellors JW, Munoz A, Giorgi JV, Margolick JB, Tassoni CJ, Gupta P, et al. Plasma viral load and CD4+ lymphocytes as prognostic markers of HIV-1 infection. Ann Intern Med. 1997;126(12):946–54.

12. Mellors JW, Margolick JB, Phair JP, Rinaldo CR, Detels R, Jacobson LP, et al. Prognostic value of HIV-1 RNA, CD4 cell count, and CD4 Cell count slope for progression to AIDS and death in untreated HIV-1 infection. JAMA. 2007;297(21):2349–50.

13. Korenromp EL, Williams BG, Schmid GP, Dye C. Clinical prognostic value of RNA viral load and CD4 cell counts during untreated HIV-1 infection--a quantitative review. PLoS One. 2009;4(6):e5950.

14. Herbeck JT, Muller V, Maust BS, Ledergerber B, Torti C, Di Giambenedetto S, et al. Is the virulence of HIV changing? A meta-analysis of trends in prognostic markers of HIV disease progression and transmission. AIDS. 2012;26(2):193–205.

15. Pilcher CD, Tien HC, Eron JJ, Jr., Vernazza PL, Leu SY, Stewart PW, et al. Brief but Efficient: Acute HIV Infection and the Sexual Transmission of HIV. J Infect Dis. 2004;189(10):1785–92.

16. Gray RH, Wawer MJ, Brookmeyer R, Sewankambo NK, Serwadda D, Wabwire-Mangen F, et al. Probability of HIV-1 transmission per coital act in monogamous, heterosexual, HIV-1-discordant couples in Rakai, Uganda. Lancet. 2001;357(9263):1149–53.

17. Quinn TC, Wawer MJ, Sewankambo N, Serwadda D, Li C, Wabwire-Mangen F, et al. Viral load and heterosexual transmission of human immunodeficiency virus type 1. Rakai Project Study Group. N Engl J Med. 2000;342(13):921–9.

18. Fideli US, Allen SA, Musonda R, Trask S, Hahn BH, Weiss H, et al. Virologic and immunologic determinants of heterosexual transmission of human immunodeficiency virus type 1 in Africa. AIDS Research and Human Retroviruses. 2001;17(10):901–10.

19. Baeten JM, Kahle E, Lingappa JR, Coombs RW, Delany-Moretlwe S, Nakku-Joloba E, et al. Genital HIV-1 RNA Predicts Risk of Heterosexual HIV-1 Transmission. Sci Transl Med. 2011;3(77):77ra29.

20. Hecht FM, Hartogensis W, Bragg L, Bacchetti P, Atchison R, Grant R, et al. HIV RNA level in early infection is predicted by viral load in the transmission source. AIDS. 2010;24(7):941–5.

21. Alizon S, von Wyl V, Stadler T, Kouyos RD, Yerly S, Hirschel B, et al. Phylogenetic approach reveals that virus genotype largely determines HIV set-point viral load. PLoS Pathog. 2010;6(9).

22. Hollingsworth TD, Laeyendecker O, Shirreff G, Donnelly CA, Serwadda D, Wawer MJ, et al. HIV-1 transmitting couples have similar viral load set-points in Rakai, Uganda. PLoS Pathog. 2010;6(5):e1000876.

23. van der Kuyl AC, Jurriaans S, Pollakis G, Bakker M, Cornelissen M. HIV RNA levels in transmission sources only weakly predict plasma viral load in recipients. AIDS. 2010;24(10):1607–8.

24. Fraser C, Lythgoe K, Leventhal GE, Shirreff G, Hollingsworth TD, Alizon S, et al. Virulence and pathogenesis of HIV-1 infection: an evolutionary perspective. Science. 2014;343(6177):1243727.

25. Payne R, Muenchhoff M, Mann J, Roberts HE, Matthews P, Adland E, et al. Impact of HLA-driven HIV adaptation on virulence in populations of high HIV seroprevalence. Proc Natl Acad Sci USA. 2014;111(50):E5393–400.

26. Eaton JW, Hallett TB. Why the proportion of transmission during early-stage HIV infection does not predict the long-term impact of treatment on HIV incidence. Proc Natl Acad Sci U S A. 2014;111(45):16202–7.

27. Eaton JW, Johnson LF, Salomon JA, Barnighausen T, Bendavid E, Bershteyn A, et al. HIV treatment as prevention: systematic comparison of mathematical models of the potential impact of antiretroviral therapy on HIV incidence in South Africa. PLoS Med. 2012;9(7):e1001245.

28. Lingappa JR, Hughes JP, Wang RS, Baeten JM, Celum C, Gray GE, et al. Estimating the impact of plasma HIV-1 RNA reductions on heterosexual HIV-1 transmission risk. PLoS One. 2010;5(9):e12598.

29. Hughes JP, Baeten JM, Lingappa JR, Magaret AS, Wald A, de Bruyn G, et al. Determinants of per-coital-act HIV-1 infectivity among African HIV-1-serodiscordant couples. J Infect Dis. 2012;205(3):358–65.

30. Gupta SB, Jacobson LP, Margolick JB, Rinaldo CR, Phair JP, Jamieson BD, et al. Estimating the Benefit of an HIV-1 Vaccine That Reduces Viral Load Set Point. J Infect Dis. 2007;195:546–50.

31. Pantazis N. Temporal trends in prognostic markers of HIV-1 virulence and transmissibility. An observational cohort study. Lancet HIV. 2014.

32. Siedner MJ, Ng CK, Bassett IV, Katz IT, Bangsberg DR, Tsai AC. Trends in CD4 count at presentation to care and treatment initiation in sub-Saharan Africa, 2002–2013: a meta-analysis. Clin Infect Dis. 2015;60(7):1120–7.

33. Porco TC, Lloyd-Smith JO, Gross KL, Galvani AP. The effect of treatment on pathogen virulence. J Theor Biol. 2005;233(1):91–102.

34. Roberts HE, Goulder PJ, McLean AR. The impact of antiretroviral therapy on population-level virulence evolution of HIV-1. J R Soc Interface. 2015;12(113).

35. Schacker TW, Hughes JP, Shea T, Coombs RW, Corey L. Biological and virologic characteristics of primary HIV infection. Ann Intern Med. 1998;128(8):613–20.

36. Bonhoeffer S, Funk GA, Gunthard HF, Fischer M, Muller V. Glancing behind virus load variation in HIV-1 infection. Trends Microbiol. 2003;11(11):499–504.

37. Geskus RB, Prins M, Hubert JB, Miedema F, Berkhout B, Rouzioux C, et al. The HIV RNA setpoint theory revisited. Retrovirology. 2007;4:65.

38. Cori A, Pickles M, van Sighem A, Gras L, Bezemer D, Reiss P, et al. CD4 cell dynamics in untreated HIV-1 infection: overall rates, and effects of age, viral load, gender and calendar time. AIDS. 2015;In press.

39. Health SADO. The 2010 National Antenatal Sentinel HIV and Syphilis Prevalence Survey in South Africa. In: Health Do, editor. Pretoria, South Africa2011.

40. Nagelkerke NJ, Arora P, Jha P, Williams B, McKinnon L, de Vlas SJ. The rise and fall of HIV in high-prevalence countries: a challenge for mathematical modeling. PLoS Comput Biol. 2014;10(3):e1003459.

41. Funk S, Bansal S, Bauch CT, Eames KT, Edmunds WJ, Galvani AP, et al. Nine challenges in incorporating the dynamics of behaviour in infectious diseases models. Epidemics. 2015;10:21–5.

42. Yerly S, von Wyl V, Lederberger B, Boni J, Schupbach J, Burgisser P, et al. Transmission of HIV-1 drug resistance in Switzerland: a 10-year molecular epidemiological survey. AIDS. 2007;21(16):2223–9.

43. Rhee SY, Blanco JL, Jordan MR, Taylor J, Lemey P, et al. Geographic and temporal trends in the molecular epidemiology and genetic mechanisms of transmitted HIV-1 drug resistance: an individual-patient-and sequence-level meta-analysis. PLoS Med. 2015;12(4):e1001810.

44. Yang WL, Kouyos RD, Boni J, Yerly S, Klimkait T, Aubert V, et al. Persistence of transmitted HIV-1 drug resistance mutations associated with fitness costs and viral genetic backgrounds. PLoS Pathog. 2015;11(3):e1004722.

45. Castro H, Pillay D, Cane P, Asboe D, Cambiano V, Phillips A, et al. Persistence of HIV-1 transmitted drug resistance mutations. J Infect Dis. 2013;208(9):1459–63.

